# Massively parallel CRISPRi assays reveal concealed thermodynamic determinants of dCas12a binding

**DOI:** 10.1101/777565

**Authors:** David A. Specht, Yasu Xu, Guillaume Lambert

## Abstract

The versatility of CRISPR-Cas endonucleases as a tool for biomedical research has lead to diverse applications in gene editing, programmable transcriptional control, and nucleic acid detection. Most CRISPR-Cas systems, however, suffer from off-target effects and unpredictable non-specific binding that negatively impact their reliability and broader applicability. To better evaluate the impact of mismatches on DNA target recognition and binding, we develop a massively parallel CRISPR interference (CRISPRi) assay to measure the binding energy between tens of thousands of CRISPR RNA (crRNA) and target DNA sequences. By developing a general thermodynamic model of CRISPR-Cas binding dynamics, our results unravel a comprehensive map of the energetic landscape of *Francisella novicida* Cas12a (FnCas12a) as it searches for its DNA target. Our results reveal concealed thermodynamic factors affecting FnCas12a DNA binding which should guide the design and optimization of crRNA that limit off-target effects, including the crucial role of an extended PAM sequence and the impact of the specific base composition of crRNA-DNA mismatches. Our generalizable approach should also provide a mechanistic understanding of target recognition and DNA binding when applied to other CRISPR-Cas systems.

## INTRODUCTION

Clustered regularly interspaced short palindromic repeats (CRISPR) and its associated genes are part of an adaptive immunity system used to combat phage infections in bacteria and archaea [1]. The system consists of two main components: a CRISPR array, which contains repetitive sequences called repeats and variable sequences called spacers, and CRISPR-associated (Cas) genes, which facilitate spacer acquisition and the destruction of foreign DNA and RNA. Mature CRISPR RNAs (crRNAs) derived from the CRISPR array can in turn program Cas nucleases to recognize and cleave DNA targets whose nucleic acid sequence is complementary with the guide portion of the crRNA and proximal to a PAM (protospacer adjacent motif) site. Due to their simple and programmable nature, the nucleases of class 2 CRISPR systems, particularly Cas9 (type II) and Cas12 (type V), have been the subject of intense research interest for the purposes of genome editing [2–4], programmable gene regulation utilizing a catalytically-dead CRISPR nuclease (dCas) [5–7], and nucleic acid detection [8, 9].

While CRISPR has already revolutionized many areas of research, from fundamental biomedical sciences to synthetic biology to disease diagnostics, a fundamental understanding of the underlying factors affecting CRISPR-Cas off-target *binding* is still lacking. This is especially important in the context of CRISPR base editors [10, 11] because off-target binding, which may not entirely correlate with DNA cleavage [12–14], needs to be reduced to a minimum level to prevent unintended base changes. While several *in silico* models [15–20] have been developed to predict the binding affinity of RNA guided CRISPR-Cas proteins using data from *in vitro* biochemical assays [21–24] or *in vivo* indel frequencies [12–14, 25–27], these approaches only provide empirical interpretations of CRISPR-Cas DNA binding and often fail to yield a conceptual understanding of the underlying factors involved in CRISPR-Cas binding. Furthermore, it can be difficult to extract quantitative binding affinity measurements from *in vivo* indel frequencies due to the inherent CRISPR-Cas binding inefficiencies associated with cellular physiological factors such as cell type, chromatin state, and delivery method [28–30]. Thus, there is a critical need for fundamental models that can help unravel the sequence-dependent determinants of CRISPR-Cas target recognition and DNA binding affinity.

To elucidate determinants of CRISPR-Cas12 off-target binding, we combine a thermodynamic model of dCas12a binding with a rationally designed CRISPRi assays to map the binding energy landscape of a type V CRISPR-Cas system from *Francisella novicida* (FnCas12a) as it inspects and binds to its DNA targets. Our approach, inspired by a recent theoretical framework that employs a unified energetic analysis to predict *S. pyogenes* Cas9 (SpCas9) cleavage activity [31] and recently developed massively parallel multiplexed assays [32–35], aims to directly measure the energetic and thermodynamic determinants of CRISPR-Cas *binding*. In other words, our assays excludes sources of variation in DNA cleavage activity caused by unknown physiological factors [28–30] by only focusing on the steps *leading* to final DNA cleavage step. Furthermore, our predictive framework is not limited to FnCas12a and can be applied to any other CRISPR-Cas systems, which should in turn fa cilitate the development of predictive models of target recognition and binding efficiency for type II and type V RNA-guided CRISPR-Cas proteins.

## RESULTS

### Thermodynamic model of dCas binding

DNA cleavage by CRISPR-Cas endonucleases may be hindered by other factors [28–30] besides the specific crRNA-DNA sequence, and it is important to disentangle these effects to gain a deep understanding of off-target binding mechanisms. We thus hypothesize that the variability in indel formation observed in live cells may not entirely originate from differences in Cas12a’s cleavage activity caused by the specific crRNA-DNA sequence targeted, but also from sequence-dependent PAM attachment efficiencies and the existence of crRNA-target DNA mismatches by asking whether the steps *leading* to a ternary complex formation play a role in CRISPR-Cas off-target binding affinity.

To formalize this approach and to obtain a fundamental understanding of the energetic landscape of dCas12a as it inspects and associates with its DNA target, we developed a general thermodynamic model of CRISPR-Cas binding dynamics to determine how crRNA-DNA mismatches affect FnCas12a target recognition and binding. This model (see supplementary information and Fig. 1) is based on recent structural biology and single-molecule studies [36, 37] which revealed that DNA hydrolysis by Cas12a occurs in three discrete stages: “PAM attachment,” where FnCas12a latches onto a PAM site, “crRNA-DNA inspection,” where FnCas12a forms a partial crRNA-DNA hybrid, and “reconfiguration,” where the protein forms a ternary complex and undergoes a conformal change that exposes its catalytic residues. While the final DNA cleaving step occurs after approximately 1 minute under the conditions tested in [36], Cas12 molecules with inactivated nuclease sites remain stably bound to their DNA target for more than 500s. Hence, the reconfiguration step effectively has no detectable *off*-rate, suggesting that DNA cleavage may be inevitable (given enough time) once Cas12a has reached this stably-bound ternary state. The same stability has also been observed in single-molecule Cas9 experiments [35].

**FIG. 1.**
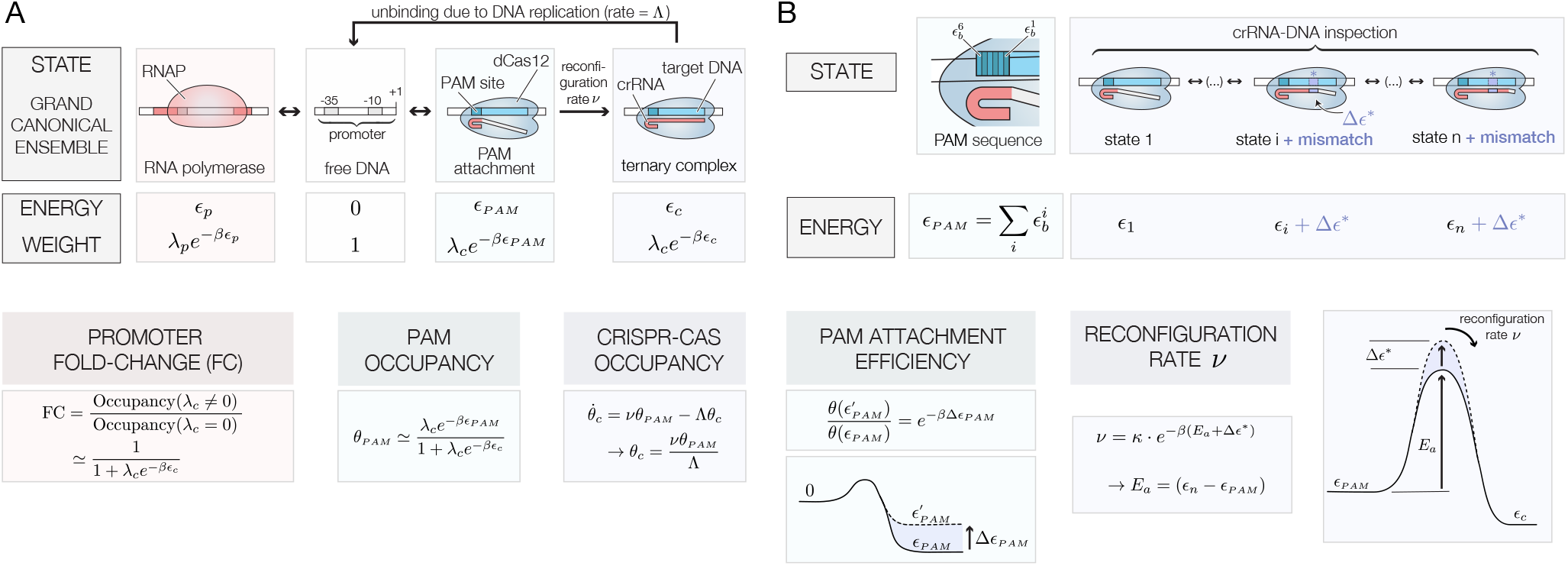
Thermodynamic model used to describe a nuclease-dead Cas12 endonuclease’s “PAM attachment,” “crRNA-DNA inspection,” and “reconfiguration” steps. A) Energy states, energies and Boltzmann weights of a dCas12a (*β* = *k*_*B*_*T*). The fold-change, PAM occupancy and CRISPR-Cas occupancy depends on the effective PAM energy *ϵ*_*PAM*_ and CRISPR-Cas binding energy *ϵ*_*c*_. All expressions assume the weak promoter 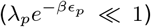 and weak PAM binding 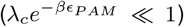 limits. B) Internal base-dependent states define a PAM specific binding energy. The specific PAM sequence dictates the relative PAM attachment efficiency between two targets. The presence of crRNA-target DNA mismatches increase the effective activation energy *E*_*a*_ and affect the effective reconfiguration rate *ν*.

Hence, our thermodynamic model describes the probability that FnCas12a loaded with a crRNA sequence will bind to a free, unobstructed target DNA sequence using the grand canonical ensemble [38–40] to derive an expression for *θ*_*c*_, the FnCas12a occupancy, which is defined as the fraction of time a DNA target will be occupied by nuclease-dead FnCas12a endonuclease. This occupancy is given by

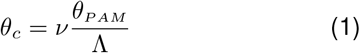

where *θ*_*PAM*_ is the PAM occupancy (the attachment probability) and *ν* is the probability that FnCas12a will form a stable ternary complex once it encounters a PAM site (the reconfiguration rate). Since DNA replication forks appear to be the only processes that can kick nuclease-dead SpCas9 off of its DNA binding site [41], we assume dCas12a unbinding occurs through a similar process–i.e. DNA duplication machinery kicks-off FnCas12a at a rate equal to Λ (the cell’s duplication rate).

We next use this approach to compare occupancies of targets that vary by a few base determinants (Fig. 1B). In this framework, the propensity of a given crRNA to target to bind to an off-target DNA region compared with its intended target is simply given by the different energetic contributions of that specific off-target location. For instance, two identical DNA targets that possess different PAM sequences have effective binding energies that differ by Δ*ϵ*_*PAM*_, which in turn translates in a reduction of the attachment probability by a factor equal to 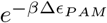 (the Boltzmann factor). Similarly, the presence of mismatches may alter the crRNA-DNA duplex energy by Δ*ϵ*\*, which in turn also yields a *e*^−*βΔϵ*\*^ change in relative binding probabilities. Hence, the *relative* binding affinity between two targets that have different PAM sites, or between an intended target and an off-target candidate, is simply given by the binding sites’ Boltzmann weight

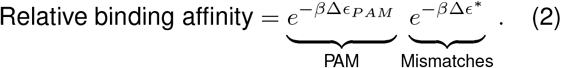

Our framework shares similarities with the uCRISPR model recently developed by Zhang *et al.* to describe SpCas9 cleavage activity [31]. However, instead of testing our model using *in vivo* indel measurements performed in human cells (which can be imprecise due to cellular physiological factors [28–30]), we use a massively parallel CRISPRi assay to directly measure the sequence-specific PAM binding energies and the energetic costs associated with crRNA-DNA mismatches in *E. coli* bacteria.

### Context dependence of FnCas12a CRISPR interference

In order to test our thermodynamic model and further explore FnCas12a target binding in *E. coli*, we developed a highly compact, 175bp-long genetic inverter inserted into a low-copy number plasmid (pSC101) containing a catalytically-dead nuclease FnCas12a (Fig. 2A, inset). The inverter element consists of a constitutive promoter driving the expression of a crRNA followed by two rho-independent terminators. Located immediately downstream of two terminators is the output promoter, which either contains a built-in PAM site within the promoter or after the promoter’s +1 location.

**FIG. 2.**
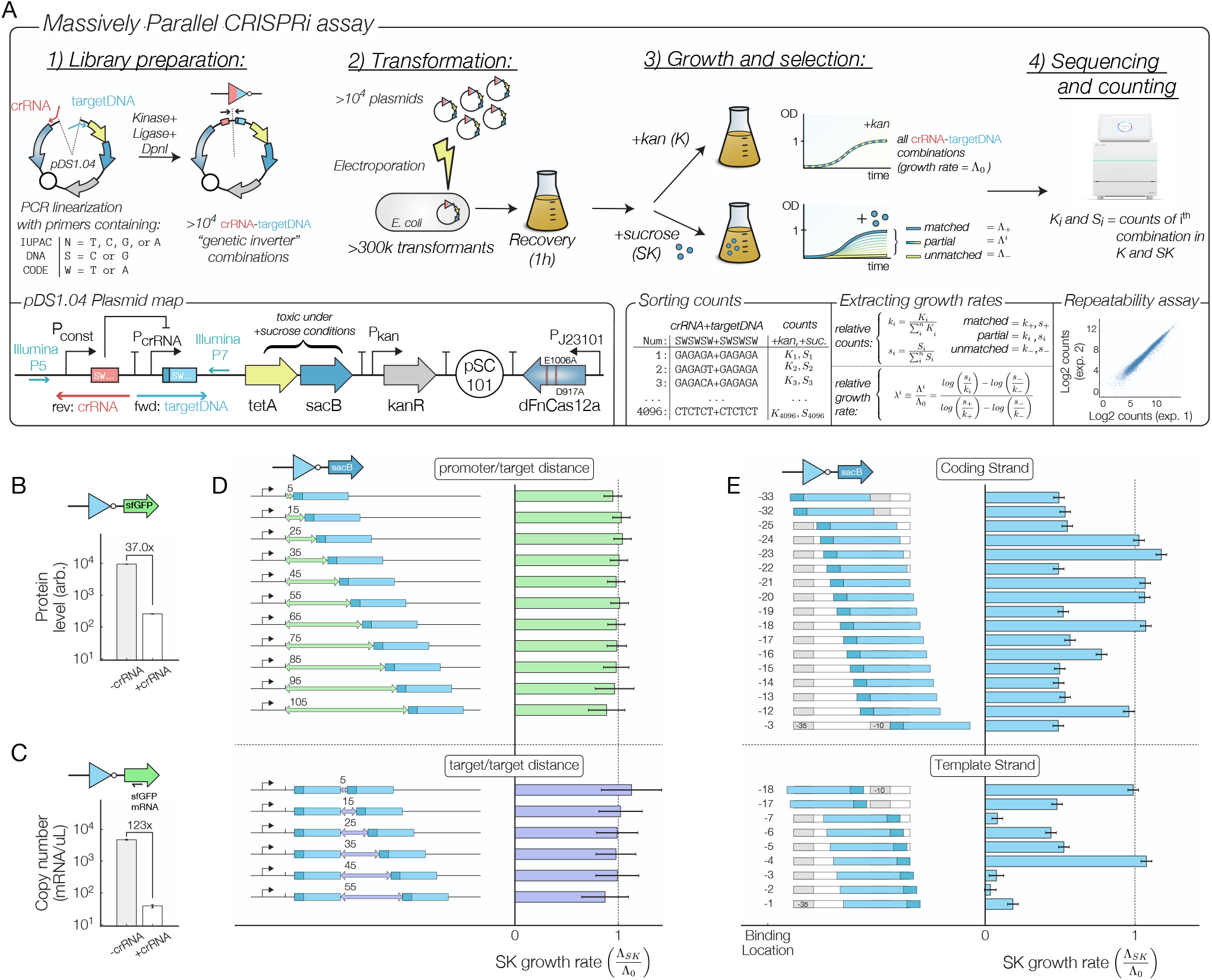
A) Experimental workflow. More than 10^4^ different crRNA-target DNA combinations are assembled in parallel using PCR primers containing degenerate IUPAC DNA codes (e.g. S, W, N). The ability of each construct to repress a tetA-sacB cassette is measured by comparing growth rates under control (K) and sucrose (SK) conditions. While the library construction is prone to some biases during the amplification and sequencing steps, a high level of repeatability is observed between experiments that started with the same assembled library (lower right). B) Protein and C) RNA level fold-change for a genetic inverter diving sfGFP expression. D) Growth rate under sucrose conditions when one or two DNA targets are located after the +1 promoter location and when E) the DNA target overlaps with the −35 and −10 regions of promoter. Λ_0_ = growth rate under control (K) conditions. Error bars are calculated using a LOESS fit [42, 43] of the mean/variance relationship between experimental replicates of the fold change.

We first sought to investigate effectiveness of Cas12a-mediated CRISPRi by measuring protein and mRNA levels of a simple inverter driving sfGFP expression. The inverter constitutively expresses a crRNA targeting a DNA binding region located at the promoter’s −19 position. Fluorescence levels for constructs containing a crRNA were 24.3 times lower than those without a crRNA (Fig. 2B) and mRNA transcript levels measured using digital droplet PCR resulted in a 123-fold reduction in mRNA transcript levels when a crRNA is expressed (Fig. 2C). Both of these results confirm that FnCas12a can repress RNA transcription [7].

Next, we tested how dCas12a interferes with RNA transcription under various configurations (Figs. 2D-E) by placing a library of up to several thousands simple inverter constructs in front of a tetA-sacB cassette. Since sacB is counterselectable genetic markers in the presence of sucrose [44] (see Fig. S2), the genetic inverters that efficiently repress RNA transcription will be enriched in the population when grown under sucrose conditions (SK). Thus, we can evaluate the ability of a RNA-guided FnCas12a to prevent transcription by comparing the number of times each construct is present in the whole population for control (K) and SK conditions using the MiSeq or iSeq100 platform from Illumina. The relative change in the population fraction is then used to find the effective growth rate Λ of every construct in each condition. While selection experiments are also performed under tetracycline-selective (TK) media, the counterselection experiment (SK) yields more useful information because the binding affinity and the dCas12 promoter occupancy is directly related to each construct’s growth rate (see supplementary materials for a complete description of this method).

Fig. 2D (top) and S3 shows that CRISPRi occurs efficiently when the FnCas12a target is located after the output promoter’s +1 transcription initiation site because the growth rate under SK conditions is close to its maximum value (Λ_0_) regardless of the location of the DNA binding site. Interestingly, while interference measurements performed using SpCas9 (a type II CRISPR-Cas nuclease) revealed that a second binding site results in suppressive combinatorial effects that multiplicatively increases CRISPRi efficiency [5], the existence of a second PAM+target DNA sequence does not improve dCas12 CRISPRi efficiency beyond what is achieved by a single target (Figs. 2D, bottom and S3).

Next, we tested FnCas12’s ability to interfere with RNA transcription initiation by introducing a PAM+target DNA sequence within the promoter sequence. In particular, we tested several inverter constructs whose PAM+target DNA sequence was located at different positions within the promoter’s −35 and −1 location, testing both the coding and template strands without altering conserved promoter regions (Fig. 2E). Our results show that CRISPR interference through promoter occlusion is efficient for most targets on both the coding and template strands, although the effective repression rate is more variable than what has been reported for CRISPR-Cas9 in terference [6]. Growth under SK conditions is also lowest when the target DNA is located on the pro-moter’s template strand at locations −1, −2, −3, and −7 with respect to the transcription initiation site, which suggests that RNA:DNA hybrids on the non-template strand display a decreased effectiveness in preventing RNA transcription initiation.

### FnCas12a binding energies depend on an extended PAM sequence

Having demonstrated the validity of our massively parallel CRISPRi assay to test multiple genetic inverter combinations, we next investigated the impact of a PAM sequence on the binding affinity of dCas12. We first tested the sequence determinant of the PAM attachment step using an oligo pool containing a degenerate 5’-NNNNNN-3’ motif for a target DNA sequence located at the promoter’s −19 position (Fig. 3A) targeted by a single crRNA (target DNA sequence=CAGTCAGTAAAATGCAGTCA). Since previous work has shown that the PAM motif required for FnCas12a DNA cleavage is TTV [3], we nevertheless tested all sequences containing up to six bases of upstream context using 4,096 PAM site variants in a single experiment. These extra bases turn out to be very important: Fig. 3B shows that while TTV is a suitable PAM site, its attachment efficiency is lower than an extended TTTV PAM site (Fig. 3B). In both individual and aggregate measurements, we observe that DNA binding to a DNA target proximal to a TTTV PAM site is 2.8 times more efficient than a TTV PAM site (Fig. 3C). This result is also confirmed by the bias towards TTTV PAM sites in the information content (Fig 3D) and the base-specific probability density in SK conditions (Fig. 3E).

**FIG. 3.**
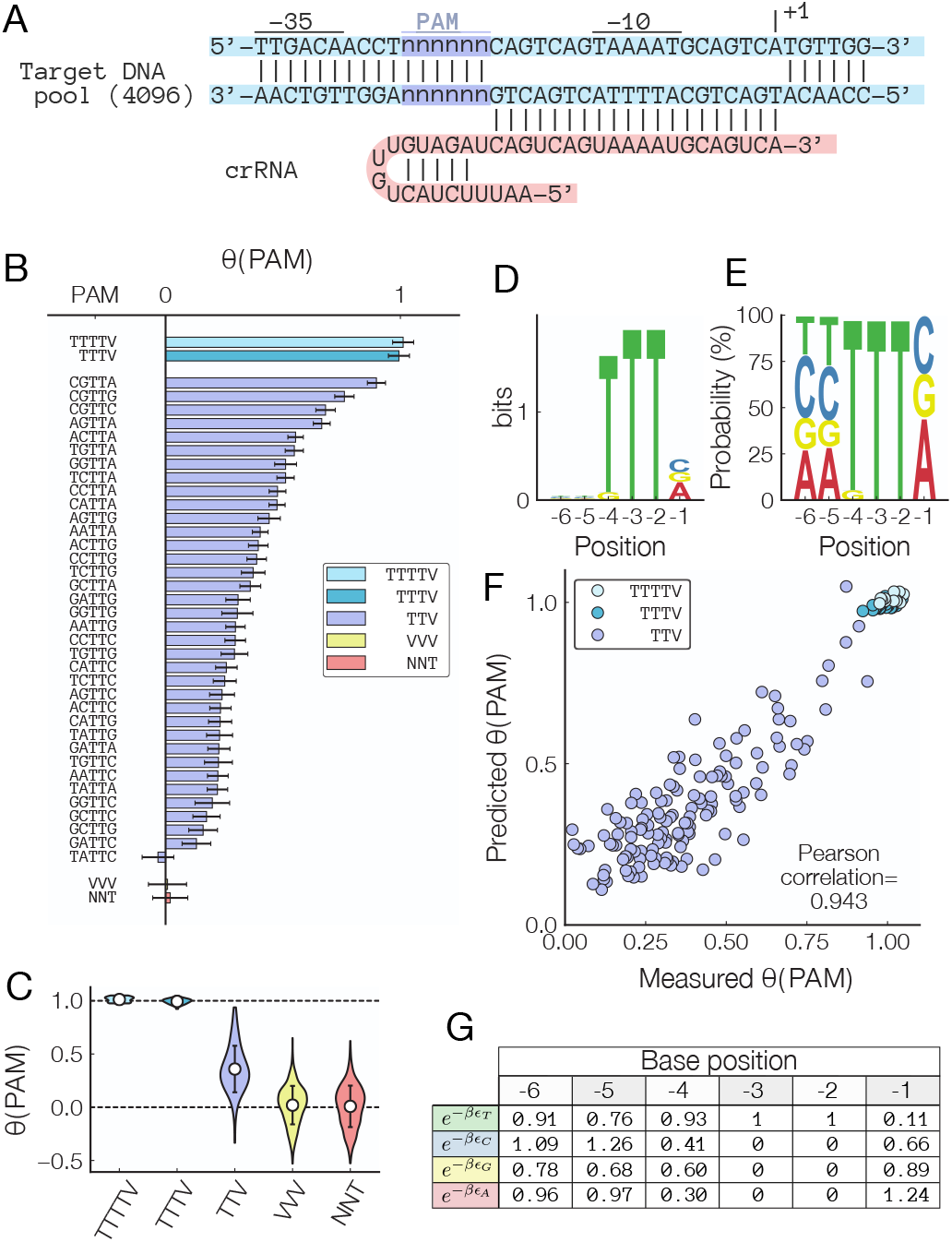
A) Sequence of the PAM site occupancy library. B) Measured PAM occupancies for all 6-base PAM sites. Error bars = aggregated LOESS fit of the mean/variance relation-ship between experimental replicates of the fold change. C) Aggregated PAM site occupancies. Error bars = std. dev. D) Bit content and E) probability density of the SK selected PAM site libraries. F) Predicted *θ*(*PAM*) using the base-dependent binding energy expression 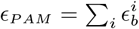 G) Fitted values for each position- and base-dependent binding energies.

Our results agree with recent work [45] which demonstrated that FnCas12a does exhibit activity in mammalian cells, but only when used with a TTTV PAM site. It is important to note that while Zetsche *et al.* [3] showed that a TTV PAM site appears to be sufficient to induce FnCas12a cleavage, it appears to be the *least efficient* motif that permits DNA binding (which could explain why FnCas12a was found to be ineffectual for mammalian cell editing using a TTV PAM site). Hence, our results suggest that PAM sites with an extended TTTV sequence should be prioritized when seeking potential FnCas12a DNA targets for CRISPRi, gene editing, nucleic-acid detection, or other applications.

Expanding on this result, we next used the measured attachment efficiencies to develop a predictive model that takes into account the full 6-base PAM site context to predict the attachment efficiency. Specifically, a natural prediction that emerges from our thermodynamics model is that the effective PAM site attachment energy is *additive*, meaning that to PAM binding energy *ϵ*_*PAM*_ of an arbitrary sequence is given by 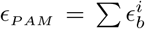, where 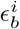 is the specific binding energy of a base of type *b*=(T,C,G,A) at location *i*=(1‥6). In this case the relative PAM binding energy between two targets 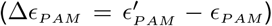 is related to the relative growth rate *λ*(*PAM*) under SK condition according to 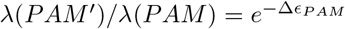.

We developed a predictive model of PAM attachment efficiency by first using an initial set of values for each 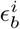 extracted from the PAM specific growth rates and optimizing the model for 1,000 additional steps to minimize the measured-predicted mean square error (see supplementary methods for details). Our model is able to accurately describe the variability in PAM attachment efficiencies observed in Fig. 3B, and its predictions for the relative PAM site occupancies *θ*_*PAM*_ agree very well with the measured attachment efficiencies (Fig. 3F, Pearson correlation = 0.943). These results suggest that PAM attachment is well described by our thermodynamic model, and the optimized energetic contribution 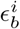 of each base *b* located at position *i* is shown in Fig. 3G. Hence, to ensure that the DNA target with the most efficient PAM site is selected when designing and optimizing a crRNA sequence for DNA binding or other gene editing application, the relative performance of each PAM sequence should be evaluated on a sequence-specific manner using the base-dependent binding energies provided in Fig. 3G.

### Off-target FnCas12a binding depends additively on mismatch energy

To better understand the impact of crRNA-DNA mismatches on dCas12 binding, we next examined how a mismatch affects the effective activation energy (Fig. 1B) that is required for FnCas12a to form a stable ternary complex. Indeed, even though a PAM site is present and dCas12 attaches itself to DNA, the additional energy associated with a crRNA-DNA mismatch can prevent DNA unzipping if insufficient homology is found. According to our model, the reconfiguration step occurs at a rate 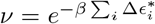, where 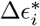 is the base-dependent energy cost associated with a single mismatche at location *i*. Thus, the location-specific energy costs associated with individual mismatches should in theory be directly obtained by measuring the reconfiguration rate *ν* of crRNA-DNA sequences that possess the same PAM sequence but with a crRNA that differs from the target DNA by one or more bases.

To test this, we used two different crRNA pools (Fig, 4A) to the mismatch-dependent reconfiguration rate *ν*. Each oligo pool consists of 4,096 different primer sequences generated by specifying degenerate DNA codes in the primer sequence (e.g. W = A or T, S = G or C), allowing us to test multiple mismatch combinations in a single experiment. Using the degenerate DNA codes S and W ensures that all crRNA sequences maintained the same GC content. In Fig. 4B, we tested the impact of “truncated” (i.e. a crRNA whose distal sequence is noncomplementary to its target DNA) and “gapped” (i.e. a crRNA whose seed sequence is noncomplementary to its target DNA) crRNAs. Consistent with other work performed in Cas12a [26, 27], our results show that optimal reconfiguration rates occur for truncated crRNAs that possess more than 15 bases of homology. Furthermore, no significant binding was detected for gapped crRNAs whose sequences that contain more than 2 mismatches.

**FIG. 4.**
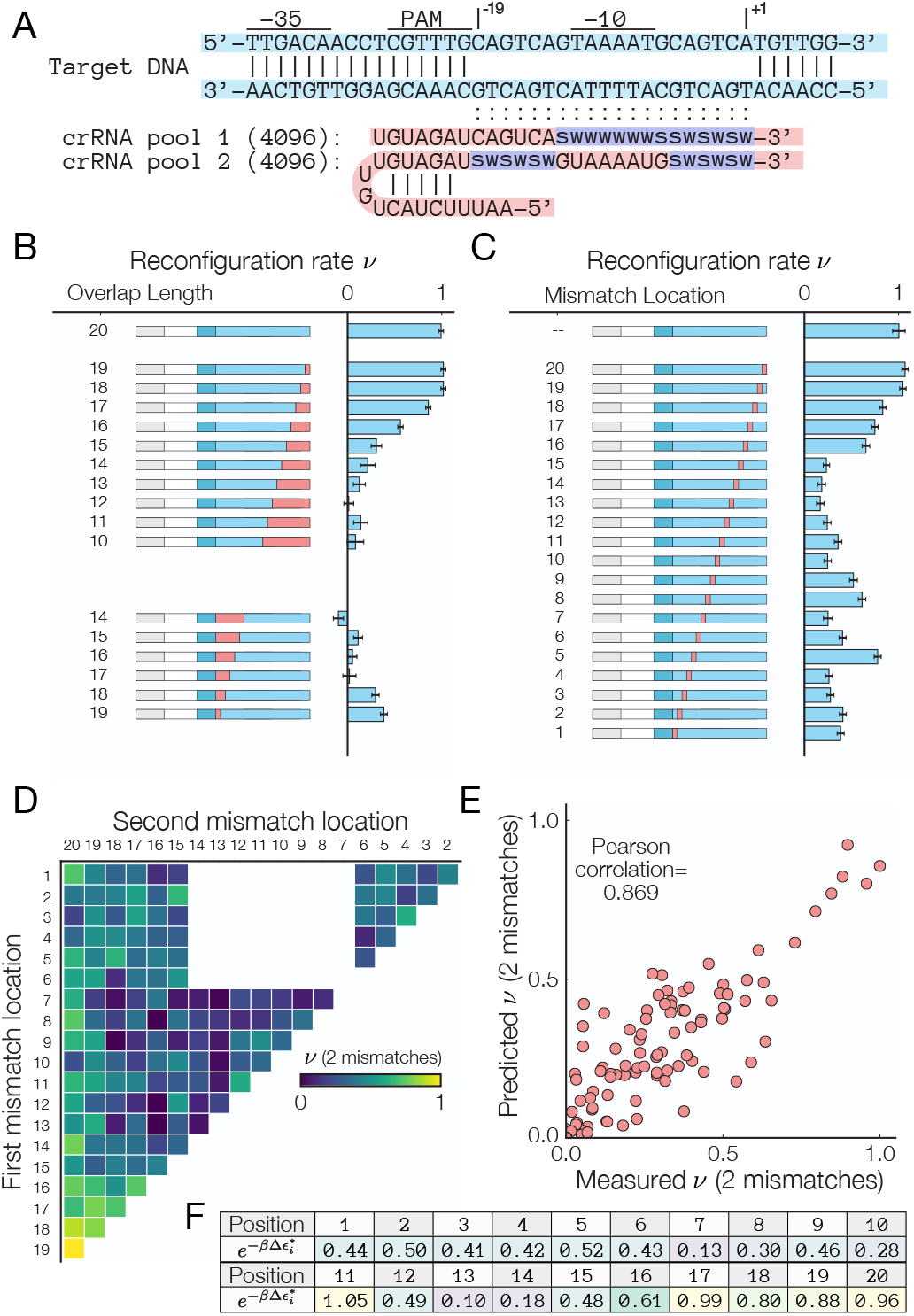
A) Sequence of the single target mismatch libraries. Reconfiguration rates for B) truncated and gapped crRNAs and for C) single mismatch crRNAs. Error bars = LOESS fit of the mean/variance relationship between experimental replicates of the fold change. D) Experimental and E) predicted reconfiguration rates for crRNA with 2 mismatches. F) Fitted values for the location-dependent binding energies for single mismatches.

Next, we measured the reconfiguration rate for crRNA containing a single mismatch (Fig. 4C). The presence of a single mismatch can decrease the configuration rate by up to 82% when the mismatch occurs in the first 17 bases of the crRNA. Consistent with prior observations by Kim *et al.* [19], the energy cost of a single mismatch does not increase monotonically with distance from the PAM site, suggesting that other contextual determinants besides position affects the reconfiguration rate *ν*. Furthermore, the presence of mismatches located in the last 3 bases of the crRNA does not impede DNA binding, confirming other works performed using *in vivo* indel measurements [26, 27] which demonstrated that crRNA-DNA mismatches negatively impact FnCas12a binding, but only in the seed and the beginning of the trunk region.

Next, we analyzed how the presence of two mismatches impacts the reconfiguration rate. Since in our model the energetic contributions 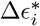 of single mismatches at location *i* are additive, we anticipated that the 2 mismatch reconfiguration rate is related to the single mismatch energies according to 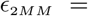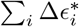. To test this, we developed a predictive model that uses the single-base mismatch energies to predict *ν*_2*MM*_. Fig. 4D shows the experimentally measured, location-dependent reconfiguration rate *ν*_2*MM*_. Using an approach similar to the one used to predict PAM attachment efficiencies, we derived baseline values for the location-dependent binding energy. While the initial Pearson correlation between the predicted and baseline energy values was initially fairly low (P=0.769), the predicted values for the two mismatch reconfiguration rate *ν*_2*MM*_ agree very well with the measured rates after the 1,000 optimization steps (P=0.869, Fig. 4E). Our results confirm that the energetic impacts of individual mismatches are additive, and location-dependent binding energy costs reported in Fig. 4F should be incorporated to models that aim to predict off-target binding.

### High throughput cross-talk assays reveal position- and nucleotide-specific energy costs

We next asked how both crRNA and DNA variations in the first six bases of the PAM-proximal seed region affected the reconfiguration rate *ν*. We performed multiplexed CRISPRi assays using two oligo pools, each containing 128 different sequences, to test the pairing between all possible crRNA-DNA sequences of the form SWSWSW or WSWSWS in a single step. Once again, those pairings were chosen to maintain all crRNA-DNA sequences at a fixed GC content. This approach covers a large combinatorial space between the spacer-target sequences and produces a comprehensive cross-talk map between 16,384 possible crRNA-DNA combinations (Fig. 5A). While we also performed the same analysis on the crRNA-DNA “trunk” region (Fig. S6), only the SW quadrant of the seed region is shown in Fig. 5B (see Figs. S4–S6 for the full cross-talk maps).

**FIG. 5.**
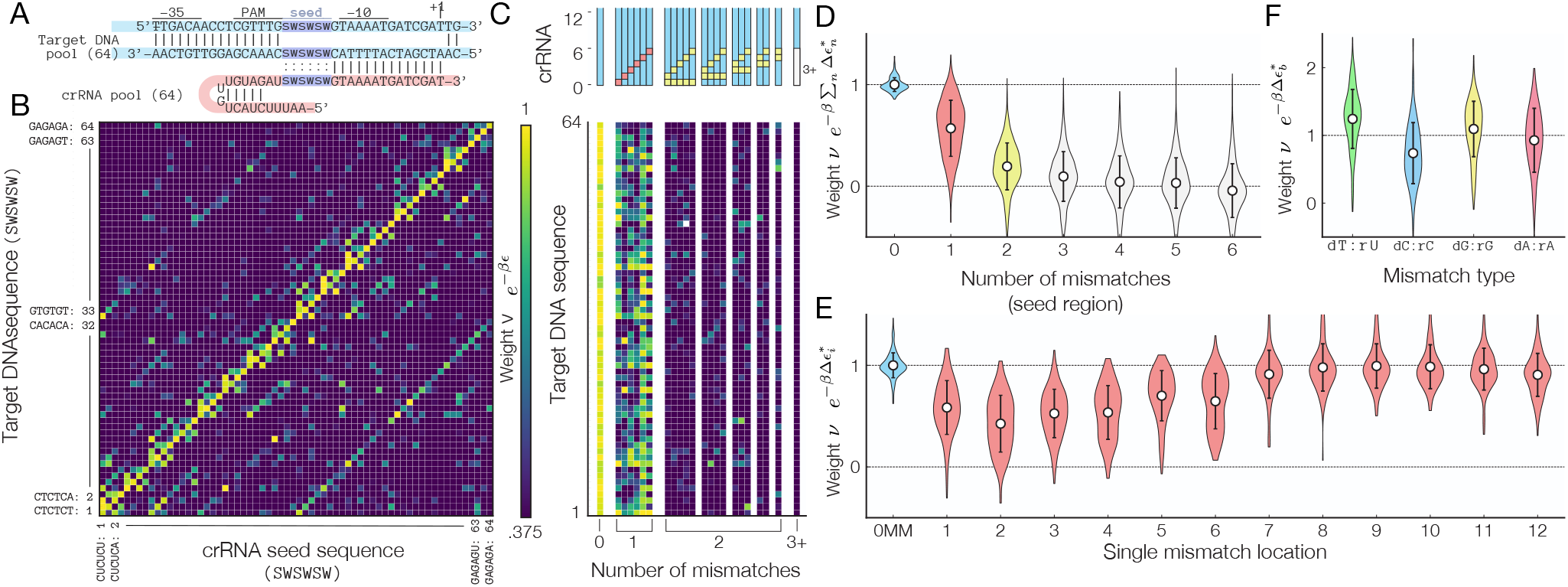
A) Sequence of the multiplexed mismatch assays for Cas12’s seed region. B) Cross-talk map of the reconfiguration rate for (SWSWSW)×(SWSWSW) data subset (full dataset is shown in Fig. S4). Note that the off-diagonal elements represent crRNA-DNA target that differ by a single mismatch. C) Target DNA dependent reconfiguration rate for SWSWSW sequences containing 0, 1, 2, or 3+ mismatches. D) Aggregated *ν* for crRNA-target DNA combinations containing between 1 to 6 mismatches. E) Aggregated *ν* for crRNA-target DNA combinations containing a single mismatches. F) The reconfiguration rate *ν* for base-specific mismatches. Note that dT:rU and dG:rG mismatches are tolerated at a higher level than dC:rC and dA:rA mismatches. Error bars = std.dev.

The cross-talk maps show that fully matching crRNA-DNA sequences (i.e. those along the main diagonal of Fig. 5B and in the first column in Fig. 5C) have the highest *ν*. Interestingly, the reconfiguration rate *ν* for all fully-matched crRNA-DNA targets fall within a very narrow range of 1.00 ± 0.06 (mean ± std.dev.), suggesting that the specific base composition of the seed region does not have a large impact on DNA binding. This contrasts with *in vivo* multiplexed DNA cleavage assays for Cas12a variants that *do* show significant sequence dependence on cleavage activity [15, 19, 20]. In addition, while SpCas9 binding and cleavage activity has different sequence specificities [12–14], we do not observe significant discrepancies between the binding and cleavage assays performed using catalytically-active FnCas12a nuclease (Fig. S7). Hence, we believe our approach may provide a more accurate representation of dCas12a’s binding energy landscape because our approach excludes any source of variation caused by unknown cellular physiological factor by only investigating a small but comprehensive portion of all possible crRNA-target DNA sequences that possess the same GC content.

To further understand how single mismatches affect the reconfiguration rate, we considered how *ν* varies as a function of the number and location of mismatches present. First, we show in Fig. 5D that no significant binding observed for sequences containing more than 4 mismatches in the seed region. Our analysis, however, reveals that formation of a stable ternary complex does occur in the presence of 1, 2 or 3 mismatches (P = 1 × 10^−232^, 8 × 10^−94^, and 1 × 10^−15^, respectively; null-hypothesis=no binding will occur for 1, 2, of 3 mismatches). It is important to note that by performing aggregate measurement across thousands of crRNA and DNA sequences, our results confers a much stronger statistical predictive power than other assays that only test a limited number of crRNA-DNA partners. In addition, we also show in Fig. 5E that mismatches have the greatest impact when located within the first 6 bases of the seed region. Sensitivity to a mismatch decreases with distance from the PAM site, and mismatches located in the trunk region (bases 6-12) only minimally impact DNA binding.

We next considered whether the type of mismatch affects *ν* in Fig. 5F. Surprisingly, we find that single crRNA-DNA mismatches of the form dC:rC decrease *ν* by an additional 26% on average. In contrast, dT:rU and dG:rG mismatches are tolerated and increase the reconfiguration rate by 9.5% and 24% compared to all types of single-base mismatch, respectively. This effect can be visualized in Fig. 5B, where off-diagonal elements that correspond to a single mismatch in the sixth location are more prominent in the lower right quadrant than those in the upper left quadrant (the upper left quadrant corresponds to a dC:rC mismatch while the lower right corresponds to dG:rG mismatches). Insensitivity to wobble-transition mismatch has been previously reported in SpCas9 [21, 46] and AsCas12a [19], but other work in AsCas12a found no significant effect due to a transversion mismatch [19], suggesting tolerance to transversion mismatches may be unique to FnCas12a.

## DISCUSSION

We have established that massively parallel CRISPRi assays, with their ability to rapidly measure thousands of different crRNA-target DNA variants in parallel, are a viable method to assess dCas12 binding efficiencies. Our results reveal the fundamental relationship between crRNA-DNA interactions and the underlying energy landscape that dictates binding behavior of dCas12. One major outcome of this study is that binding of DNA by CRISPR-Cas12a endonuclease does not strongly depend on the specific crRNA sequence used (at least within the set of tested sequences which were kept at 50% GC content). Rather, variance in DNA binding affinities depends on the PAM sequence, the presence of mismatches, and the type of mismatch present. Indeed, the propensity of identical DNA targets to be recognized by a CRISPR-Cas nuclease matching crRNA may be significantly different depending on their respective 6-base PAM sequence. Similarly, the absolute number of mismatches in the seed region of a crRNA-DNA hybrid is more important than their specific location, and mismatches that occur in the distal region of a crRNA (i.e. after base 17) do not significantly affect binding affinity. Our results also show that dT:rU and dG:rG mismatches are tolerated at a higher level than dA:rA and dC:rC mismatch.

Beyond that, the power of our approach also resides in our ability to use a parameter-free statistical mechanics framework to extract thermodynamic determinants of dCas12a binding. Importantly, our results are not specific to nuclease-dead CRISPR-Cas endonucleases –we confirm in Fig. S7 that the same behavior is observed for *catalytically-active* Cas12a nuclease– and our approach should foster the development of predictive, parameter-free biophysical models of on- and off-target binding affinities and DNA cleavage activities. In addition, because CRISPR-Cas systems are very common amongst prokaryotes [1], there is a need for the rapid and efficient characterization of newly-sequenced CRISPR-Cas systems that may display enhanced target differentiation capabilities or alternative PAM site compositions. We anticipate that this method will also provide a mechanistic understanding of the thermodynamic determinants of DNA target recognition and binding affinities in uncharacterized CRISPR-Cas endonucleases and other nucleic-acid binding enzymes.

Because our method is applicable to both the catalytically active and dead versions of the nuclease, it should also lead to improvements in a vast range of CRISPR applications, including *in vivo* gene editing, programmable repression, and nucleic acid detection. Our multiplexed approach is particularly applicable to the advancement of dCas-based gene circuit elements, which can be been used to create complex circuits that behave orthogonally, operating independently without crosstalk [47–51]. Furthermore, our approach can expedite the rational design of enhanced CRISPR nucleases and facilitate the development of CRISPR-Cas variants with greater specificity, improved proofreading capabilities, or increased activities [52–57].

## AUTHOR CONTRIBUTIONS

Conceptualization, G.L. and D.A.S.; Methodology, D.A.S. and G.L.; Formal Analysis, D.A.S. and G.L.; Investigation, D.A.S., Y.X., and G.L.; Writing – Original Draft, G.L. and D.A.S.; Writing – Review & Editing, G.L., D.A.S., and Y.X.; Supervision, G.L.;

## DECLARATION OF INTERESTS

The authors declare no competing interests.

## I. EXPERIMENTAL METHODS

### Assembly of the CRISPR-Cas12a plasmid back-bone

Unless indicated otherwise, all experiments were conducted using a plasmid backbone which constitutively expresses dCas12a (*Francisella novicida*) and tetA/sacB. This plamid was assembled using standard Gibson assembly techniques from components sourced from several other plasmids: pY003 (pFnCpf1_delta Cas) was a gift from Feng Zhang (Addgene plasmid # 69974), pTKLP-tetA was a gift from Thomas Kuhlman (Addgene plasmid # 71325), and pKM154 was a gift from Kenan Murphy (Addgene plasmid # 13036), using a backbone derived from pUA66 [1]. FnCas12a was made to be catalytically inactive via two mutations, D917A and E1006A performed using NEB’s Q5 site-directed mutagenesis kit. The landing pad sequence needed for Illumina sequencing was inserted using an IDT gBlock gene fragment (Supplementary Table 1). The entire plasmid sequence (pDS1.04) can be found here: https://benchling.com/s/seq-I9k4w1wRsX3B3cXVzyE2.

### Design of PAM and gRNA mismatch assays

In order to test the effects of PAM and gRNA mismatches at a large scale, we created a highly compact dCas12a repressing element such that target and gRNA properties could be changed with a single site-directed mutagenesis. The sequence of this compact element can be found here: https://benchling.com/s/seq-BAWu6Ya1kAnhxugFEezi.

### Assembly of plasmid libraries

Our method of exploring CRISPR interference is predicated on the use of large, randomized oligos in order to produce many mismatch combinations via site-directed mutagenesis. Oligos for PCR-based assembly of different guide:target variants were purchased from Thermo Fisher; oligos containing randomized bases were PAGE-purified and all others were ordered as desalted oligo plates. Oligonucleotide sequences are listed in Supplemental Table 1. PAGE-purified oligos were ordered phosphorylated by the manufacturer. Unphosphorylated oligos from plates were pooled together (according to their forward-reverse directions) and phosphate groups were added using Thermo Fisher’s T4 Polynucleotide Kinase (T4 PNK).

Pooled or randomized phosphorylated oligos were used to insert multiple crRNA and target DNA combinations in a single PCR step. Likely due to the large size of the insertion, we had a significant amount of difficulty finding parameters which resulted in complete PCR products. Parameters that worked were found serendipitously and include a high molar ratio of template to primers and extremely long (15min+) extension times. PCR was done exclusively using Q5 hot start DNA polymerase from NEB.

For cloning of single constructs, ligation and phosphorylation was accomplished using the Ki-nase+Ligase+DpnI (KLD) mix from NEB’s sitedirected mutagenesis kit. In the multiplexed experiments (except when noted below), ligation was accomplished using NEB’s ElectroLigase, using 100 ng of DNA from the PCR purified using Zymo’s ZymoPURE Miniprep kit. Ligation was done according to the manufacturer’s instructions, with a 60 minute incubation time at 25°C and a 15 minute inactivation step at 65°C. Ligated product was either used immediately for transformation or frozen for future use.

The catalytically active Cas12a experiment was cloned using a library derived from the Kanamycin-selected control in the catalytically-dead experiment, since this was of known good coverage for all mismatch combinations. D917A and E1006A mutations in dCas12a in pDS1.04 were reverted using site-directed mutagenesis, and the catalytically-restored Cas12a was inserted into the linearized backbone with all 4,096 variants in lieu of the catalytically-dead CRISPR via assembly with NEB Hifi DNA assembly Master Mix.

Insertions for the promoter/target and target/target spacing experiments were done using two rounds of PCR, the first one to add a functioning inverter element and the second one to add one or two PAM+target DNA sequences. Primer sequences are listed in Supplementary Table 1.

### Electroporation of plasmid libraries

In order to achieve the transformation efficiencies required for good statistical coverage of all mismatch combinations in our multiplexed experiments, we used electroporation of our CRISPR mismatch libraries. 1 *μ*L of electroligated product was added to 25 *μ*L Lucigen Endura ElectroCompetent cells, and then electroporated at 1400V (BTX ECM399 Device). Cells were recovered in 2mL of Lucigen recovery media, as in [2]. Following the one-hour recovery, the full 2mL was transferred to 23 mL of Terrific Broth (TB) with kanamycin in a 50 mL tube. TB was made by autoclaving 23.8 g of VWR’s Terrific Broth powder with 2 mL of glycerol and 500 mL of purified water. Since the Endura cells are so densely packed, the resulting recovery product has a nonzero OD of roughly 0.3. Once the tubes reached an OD of 1.0 (approximately 8 hours at 37C, 225 rpm), each pair of tubes was combined in a flask and 1 mL of that product was used to inoculate each of the selection conditions.

### Sucrose and tetracycline selection

Inoculated selection media (100mL) were grown in 250 mL flasks (37C, 225 rpm) until they reached an OD of 1.0, then cooled to 4°C prior to plasmid extraction. Unselective media (the control condition) is TB with Kanamycin (50*μ*g/mL). Tetracycline-selective media (TK - indicating both kanamycin and tetracycline) was produced in the same way, adding tetracycline at a concentration of 10*μ*g/mL. Sucrose-selective media (SK) was produced by combining 10mL of an autoclaved sucrose premix solution (22.5 g sucrose in 37.5 mL water) with a TB premix solution such that the resulting solution contains 4.5% sucrose (w/v).

Plasmid extraction was done using Zymo’s ZymoPURE II Midiprep kit according to the manufacturer’s instructions. Plasmids were then eluted in elution buffer and stored at −20°C prior to indexing for Next-Generation Sequencing.

While Li *et al.* utilize dual sensitivity to both sucrose and fusaric acid [3], we found no selective advantage due to the use of fusaric acid, and did not utilize it beyond preliminary experiments.

### Next-Generation Sequencing and Analysis

Our method is made possible by the inclusion of sequences flanking the inverter site of interest (see the pDS1.04 sequence) to which Illumina indexing primers can bind. This allows us to lift out purely the sequences of interest using PCR, skipping most traditional library preparation steps. Indexes were added to our samples using primers from NEB’s NEBNext Multiplex Oligos for Illumina (Index Primers Set 1), using NEBNext Q5 Hot Start HiFi PCR Master Mix or NEB-Next Ultra II Q5 Master Mix.

Sequencing was either performed using Illumina’s MiSeq System from the Cornell Genomics Facility (150 bp kit, PE 2 × 75 bp) or an Illumina iSeq instrument in our own laboratory (2 × 150 bp run). Due to the extremely low complexity of these libraries, a 10% PhiX spike-in was used in both cases.

Results were analyzed using scripts written in Python, which can be made available upon request. Only reads that perfectly matched the correct design in the sequencing window were counted in the final result to calculate the relative fraction of each construct in the sequenced populations.

### Fluorescence Measurements of Protein Fold-change

In initial fluorescence measurement experiments, plasmids containing dCas12a, guide RNA sequence, and a GFP target were transformed into NEB’s 5-alpha Competent *E. coli* (High Efficiency) and recovered in SOC according to the manufacturer’s instructions. Initial assessment of repression efficacy was made by visual inspection of cells grown on LB plates. The sequences of these plasmids can be found here:

- GFP Control: https://benchling.com/s/seq-D4cjbT6qdnF7bOit9Poa
- Single Inverter: https://benchling.com/s/seq-UtdJRWJ4oUW35cptcnjZ.

Quantitative measurements of fluorescence (used to produce Fig. 2B) were made using a Synergy H1 Hybrid Multi-Mode Microplate Reader, produced by BioTek. Reported fold-change corresponds to asymptotic fold-change observed after roughly five hours of growth in 200uL TB at 37C. GFP fluorescence measurements are corrected by subtracting out the measured green emittance from cells at the same OD which entirely lack GFP.

### ddPCR Measurement of mRNA fold-change

mRNA fold-change was measured using droplet digital PCR (ddPCR) measurements. Transformed cells were grown in 20 mL TB for 12 hours at 37C and 300uL was then used for RNA extraction using Zymo’s Direct-zol RNA MiniPrep. Genomic DNA was removed using Thermo’s TURBO DNA-free Kit and 10ng of cleaned RNA was then used as a template for cDNA production, utilizing the ProtoScript II Reverse Transcriptase kit from NEB and primer RT_GFP_Rev from Supplementary Table 1.

Droplet generation was done using a QX200 Droplet Generator produced by Bio-Rad. PCR amplication was done using a C1000 Touch Thermal Cycler (Bio-Rad) utilizing EvaGreen Supermix and following the manufacturer’s instructions. Primers corresponding to the GFP target are listed in Supplementary Table 1. Results are read out on a Qx200 Droplet Reader. Data analysis from ddPCR was completed using QuantaSoft software made available by the Cornell Genomics Center.

### Measurement of cell growth as a function of sucrose and tetracycline concentration

The Synergy H1 microplate reader was used to produce growth curves for cell growth in the presence of sucrose and tetracycline. Cells with sacB (pDS1.04) were tested with varying concentrations of sucrose and cells lacking tetA were tested against varying concentrations of tetracycline. Cells were grown in 200uL TB at 37C. Growth rates reported in Supplementary Figure 2 are the result of a logistic curve fit to the optical density measurement, fixed such that each curve has a constant starting OD.

### PAM site sequence logo

Since sequencing coverage was in excess of 100x for most sequences, all sequences were still detected under SK conditions, including those that had a Λ_*i*_ = 0 growth rate. Hence, to generate the sequence logo and final base density in Fig. 3D and E that were not tainted with those Λ = 0 sequences, simulated counts 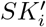 were used instead of the measured counts *SK*_*i*_. These simulated counts 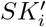 were computed from K-condition counts *K*_*i*_ according to 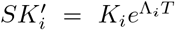, where T=17.5/Λ_0_, an arbitrary growth time, and Λ_*i*_ is the growth rate of each PAM sequence (see *Growth rate from sequencing counts* section below). Then, sequence logos were computed from *baseheight* = *f*_*b*,*i*_*R*_*i*_, where *f*_*b*,*i*_ is the relative frequency of base *b* at position *i* and *R*_*i*_ = *log*_2_(4) − ∑_*b*_(−*f*_*b*,*i*_)*log*_2_(*f*_*b*,*i*_).

### Statistical analysis and confidence interval evaluation

While it is is prohibitive to replicate next generation sequencing experiments, there are independent replicates within a single experiment with different selection conditions from which we can extract a variance as a function of the number of counts from next generation sequencing. Specifically, independent replicates are sourced from the K and TK selection conditions with 6 mismatches in the seed region, for which we expect there to be no effective repression by dCas12a.

We utilize the transformation 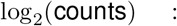 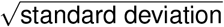 previously used by other authors for RNA seq counts [4]. This transformation is then fit using LOESS [5] via its python implementation [6]. Variance falls with the log of the number of counts (as would be expected from Poisson statistics) but then asymptotes for large counts.

### Data Availability

The raw fastq files from sequencing and data generated during this study are available at https://www.ncbi.nlm.nih.gov/Traces/study/?acc=PRJNA549693.

## II. MASSIVELY PARALLEL CRISPR INTERFERENCE ASSAY

### Growth rate from sequencing counts

Each experiment contains a total of *n* different crRNA:DNA combinations. The total number of transformant after plasmid assembly is 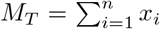, where *x*_*i*_ is the number of cells with a specific crRNA-target DNA combination *i*. If the assembly of each feature *x*_*i*_ does not depend on the underlying DNA sequence, the distribution of *x*_*i*_ will be given by a Poisson distribution with rate 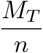.

The sample is then grown under two different selection conditions (K and SK) until it reaches an optical density OD_600_=1. The time needed for each the population to reach OD_600_=1 in each condition is given by *τ*_*K*_ and *τ*_*S*_, respectively. Plasmids are collected when each flask reaches OD=1 and the region containing the crRNA and target DNA coding sequence is amplified using NEBNext Multiplex Oligos for Illumina (Index Primers Set 1).

Each sample is then sequenced using either the Miseq or iSeq 100 platform, and the number of times each crRNA-target DNA combination *i* is present in the population after selection is denoted by *K*_*i*_, *S*_*i*_, and *T*_*i*_. Each feature *i* will grow at a rate 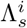 under SK selection and at a constant rate 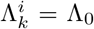 under K selection.

We define the relative fraction of each feature *i* in each condition according to:

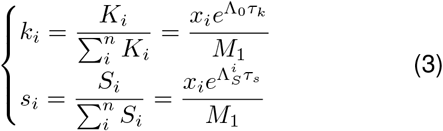

where *M*_1_ is the number of cells at the end of the experiment when the flask reaches OD=1.

To find the effective growth rate 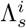 under SK-condition for all other features *i*, we can re-arrange eqn. 3 to get

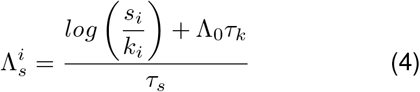

### Determining the growth time *τ*_*k*_ under K-selection

A subset of the population will not fully repress the tetA-sacB cassette and will not grow in SK conditions. This happens, for example, when the crRNA/target DNA hamming distance is 6. If we denote the population fraction under K and SK selection conditions of this non-growing subpopulation as *s*_−_ and *k*_−_, we get

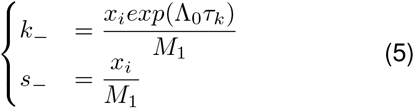

Using this, we can find the time *τ*_*k*_ cells were growing under K-selection from the ratio 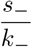 and obtain

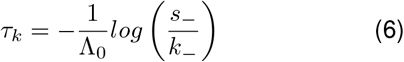

### Determining the growth time *τ*_*s*_ under SK-selection

Next, consider the subpopulation that is expected to grow at the same rate in either condition, which should occur when the crRNA/target DNA Hamming distance is zero. Using the population fractions in the K and SK conditions for this subpopulation (labeled *s*_+_ and *k*_+_, respectively), we can use Eqn. 3 to find *τ*_*s*_:

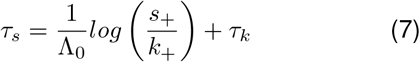

Here, we assumed that 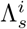 (the growth rate under SK conditions) is equal to Λ_0_ (the growth rate under K conditions) for all combinations *i* with a hamming distance of zero.

### Determining the growth rate Λ_*s*_ under SK-selection

Having derived expressions for *τ*_*k*_ and *τ*_*s*_, we can expand eqn. 4 to get an expression for the relative growth rate 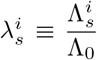 in terms of the *s*_±_ and *k*_±_ population fractions:

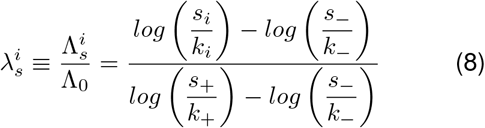

### Determining the growth rate Λ_*t*_ under TK-selection

Using a similar approach, we also derive an analogous expression of the growth rate 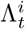 under TK-selection, assuming the *t*_+_ cells grow in TK when Hamming=6 and the *t*_−_ cells do not grow when Hamming = 0. Specifically, we get:

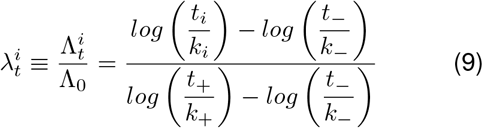

### Determining the initial population sizes x_i_

While the size of each founding population *x*_*i*_ cancels out in our analysis, we still need to find its probabilistic distribution in order to compute its expected variance from the measured number of counts. Specifically, if we consider the measured population fraction under K-selection:

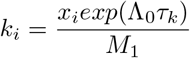

we note that since the factor 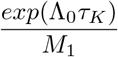 is common to all sequences, we can set *x*_*i*_ = *k*_*i*_.

## THERMODYNAMIC MODEL OF CRISPR-CAS BINDING

### Measuring dCas occupancy from the grand canonical ensemble

To measure the effective dCas occupancies *θ*_*c*_ for the tested targets, we use an auxiliary reporter system to measure the effective dCas occupancy *θ*_*c*_. Consider a simple CRISPR interference (CRISPRi) promoter architecture, where a protospacer adjacent motif (PAM) and a target DNA overlaps with the −35 or −10 consensus site of the promoter (Fig. 2). Binding of an RNA-guide CRISPR-Cas endonuclease with deactivated nuclease sites (dCas) to the promoter prevents initiation of RNA transcription by RNA polymerase (RNAP). In this scenario, the binding energy of the CRISPR-Cas protein to its target DNA site is *ϵ*_*c*_, the RNAP binding energy is *ϵ*_*p*_, the PAM site binding energy is *ϵ*_*PAM*_, and the grand partition function of this system is [7]

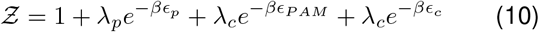

where *β* = *k*_*B*_*T*, *μ*_*p*_ and *μ*_*c*_ are the RNAP and dCas chemical potentials 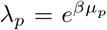 and 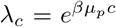 are the RNAP and dCas fugacities.

Using 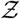, we derive an expression for the fold-change, defined as the ratio of the average number of absorbed RNAP molecules with and without repressor molecules [8], and get

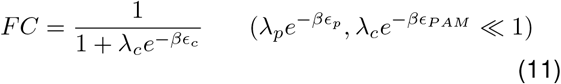

Here, we used the weak promoter limit 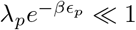 because the RNAP typically initiates RNA transcription immediately after binding to the promoter [7] and does not occupy the promoter for a long time (i.e. it binds to the promoter in a manner that appears as though its binding energy is very weak). Similarly, we used a weak PAM binding limit 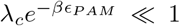 because we assume that the dCas protein typically does not remain in the PAM-bound state for a long time (approximately 0.13s according to single-molecule studies [9]) and will only transitions into a stable ternary complex if sufficient crRNA-DNA homology is found.

The average dCas occupancy is

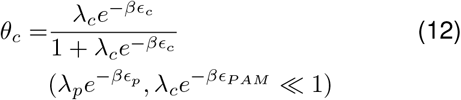

In terms of the fold-change, *θ*_*c*_ becomes

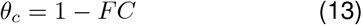

Hence, starting from the fold-change, an easy to measure quantity, we can extract the effective occupancy probability of a RNA-guided dCas protein to its DNA target. In Fig. 2B, the fold-change for a perfectly matching crRNA-DNA hybrid measured using ddPCR is 1/123, meaning that the quantity 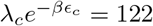 and *θ*_*c*_ = 122/123 = 99.2%.

We can also extract the average PAM site occupancy *θ*_*PAM*_ according to

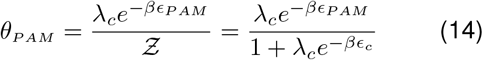

### Measuring dCas occupancy from transition state theory

The initial attachment step involves recognition of a PAM site by a crRNA-loaded Cas12 endonuclease. This recognition step depends on the specific PAM sequence, and leads to a conversion into a fully-bound state with probability *ν* if sufficient crRNA-target DNA homology is found. Then, the only way for a stably bound dCas protein to unbind its target DNA is to be destabilized by the DNA replication machinery [10]. Hence, given a PAM binding energy *ϵ*_*PAM*_, a (PAM)→(stable complex) transition probability given by the reconfiguration rate *ν*, and a growth rate Λ, we obtain

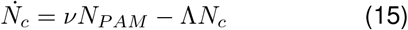

where *N*_*c*_ and *N*_*PAM*_ are the number of dCas proteins bound to their target DNA and bound to the PAM site, respectively.

At steady state, we note that *N*_*tot*_*θ*_*PAM*_ = *N*_*PAM*_ to derive an expression for *θ*_*c*_ and get

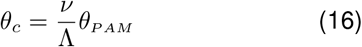

To find *θ*_*PAM*_, we can extract the sequence-specific binding energy 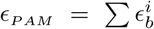 by keeping the same target DNA and measuring *θ*_*c*_ for different PAM sequences. Specifically, a PAM sequence which deviates from the canonical PAM site sequence will have a binding energy given by 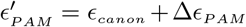, which will decrease the PAM occupancy *θ*_*PAM*_ by a factor 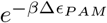 in the weak PAM binding limit.

To find *ν*, we first assume that the (PAM)→(stable complex) transition occurs in a number of *n* discrete steps, and each step *i* can only transition to either state *i* − 1 or *i* + 1. In this case, the transition rate from state 1 state *n* is simply given by:

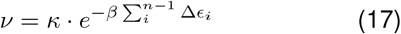

where Δ*ϵ*_*i*_ = *ϵ*_*i*+1_ − *ϵ*_*i*_ and *κ* is a pre-exponential factor assumed to be constant for all experimental conditions. However, if a crRNA-target DNA mismatch exists at location *i*, the new binding energy will be 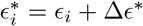 and the new rate *ν* will be given by

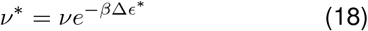

Combining equations 16 and 18, we obtain an explicit formulation of the relative binding probabilities between two targets:

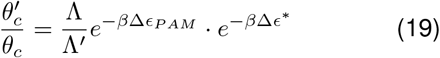

When measured in *E. coli* bacteria, both the DNA replication rate and the thermodynamic determinants of Cas12 binding will impact 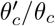. We describe in the next section how we untangle both effects using our massively parallel CRISPRi assay.

In mammalian cells, on the other hand, DNA replication rates are not affected by CRISPR-Cas binding (i.e. Λ = Λ′). In this case, only the thermodynamic determinant of Cas12 binding (i.e. the PAM attachment probability and the reconfiguration rate) will have an impact on DNA binding probabilities. Since DNA binding is also directly related to DNA cleavage activity, the relative indel frequency between two targets is thus given by

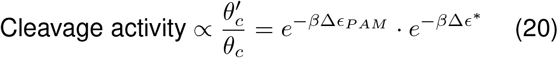

Thus, knowing the basic thermodynamic determinants of Cas12 binding can help determine the relative cleavage activity between any two DNA targets.

### dCas12 occupancies from growth rate

The tetA-sacB cassette is under the control of a dCas12 repressible promoter whose fold-change expression (Eqn. 11) is given by

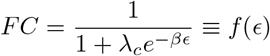

where the fugacity *λ*_*c*_ converges to the concentration of dCas12+crRNA binary complex in the [crRNA] ⨠ 1 limit and *E* is the effective binding energy of the stabilized dCas12+crRNA+DNA ternary complex.

We describe the kinetics of this system using this system of ODE equations:

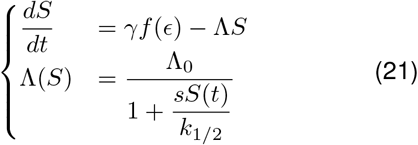

where *S* is the number of sacB molecules, *s* is the sucrose concentration, *γ* is the sacB production rate, and the growth rate Λ is given by Monod kinetics with *k*_1/2_ is the half-velocity constant and maximum growth rate Λ_0_.

At steady-state, we get:

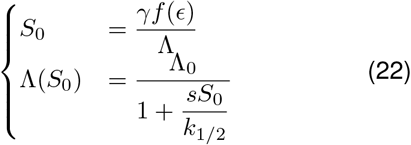

Solving for Λ in the quadratic equation generated, we obtain

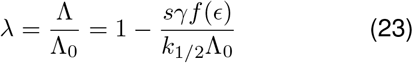

To find *k*_1/2_, we measured the growth rate of cells that constitutively express sacB as a function of sucrose concentration and get that *k*_1/2_ = *s*_1/2_*S*_*max*_, where *S*_*max*_ is the maximum sacB level produced when *f*(*ϵ*) = 1. From Fig. S2, *s*_1/2_ = 0.6% sucrose and since the experiments were performed at a sucrose concentration of 4.5%, we can rearrange Eqn. 23 to get

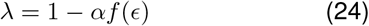

where 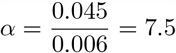.

The dCas12 occupancy *θ*_*c*_ is related to the fold-change and growth rate 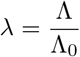 according to

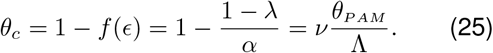

Hence, given two arbitrary crRNA-target DNA configurations, the ratio of their dCas12 occupancies is

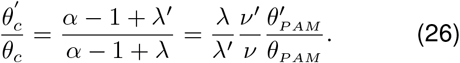

Noting that *α* ⨠ 1, we obtain

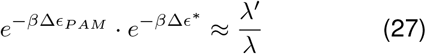

where Δϵ_*PAM*_ and Δ*ϵ*\* are the PAM and mismatch binding energy differences, respectively.

Eqn. 27 allows us to untangle the contribution of the thermodynamic determinants of Cas12 binding from growth-dependent effects due to tetA-sacB expression. As mentioned in the previous section, the thermodynamic determinants of Cas12 binding (parametrized as *ϵ*_*PAM*_ and Δ*ϵ*\*) can then be used to evaluate the relative cleavage activity and indel frequency between different DNA targets in any context (including for genomic edits in mammalian cells).

### Measuring dCas occupancy for TK selection

When a crRNA regulates the expression of tetA or targets an essential gene, the growth rate Λ depends on the amount of tetA proteins *T* and tetracycline concentration [*tet*] in the cell. If we measure the growth rate of cells that constitutively express tetA as a function of tetracycline concentration (Fig. S2), we get that the half-max growth rate occurs for [*tet*]_1/2_=0.14*μ*g/mL. This means that the experiment were carried out at a 10*μ*g/mL tetracycline concentration, any fold-change *f*(*ϵ*) smaller than 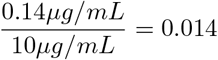 will prevent cells from growing. Therefore, even a partially-repressed tetA-sacB cassette will decrease the amount of tetA in the cell and can in turn drastically reduce the growth rate. We are therefore unable to detect differences in dCas12 binding by monitoring the growth rate alone, and we mainly use the TK growth rates as a means to confirm repression trends observed under SK selection.

## MODEL PREDICTIONS

### Fitting the PAM attachment and mismatch costs

According to Equation 20, two identical DNA targets that are flanked by different PAM sequence will have the same reconfiguration rate *ν* but different PAM attachment energies, which in turn will yield different growth rates under SK conditions. In our model, the PAM attachment energy is defined as 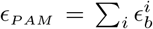 where 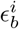, is the specific binding energy of a base of type *b*=(T,C,G,A) at location *i*=(1..6). In Fig. 3F and G, we use SK growth rates for PAM sites of the form NNNTTV to compute the position-dependent binding energies 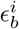.

First, we computed a *baseline* value for all 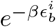 the Boltzmann weight of base *b*=(T, C, G, A) at location *i*, by averaging all the growth rates of the PAM of the form NNNTTV. Specifically,

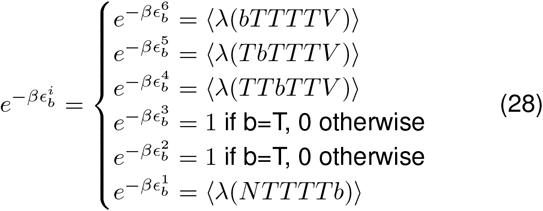

where the brackets 〈·〉 signify averages over either *V* =(C, G, A) or *N* =(T, C, G, A).

This process first yielded this unoptimized energy “matrix”:

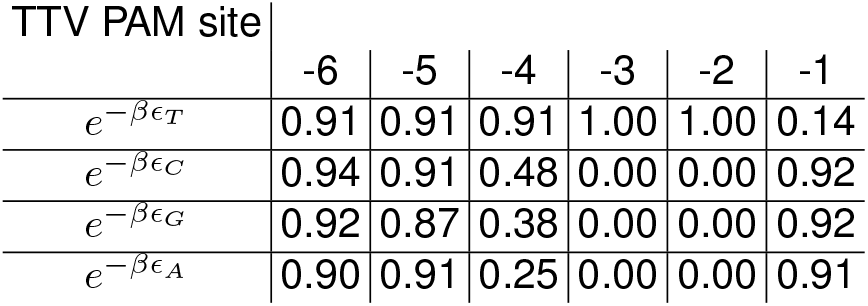

We then optimized the energy matrix for the TTV PAM sites by performing 1,000 optimization steps where we 1) added normally distributed noise 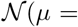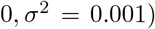 to each value of the energy matrix, 2) used the new matrix to compute new predicted values for *λ*(*PAM*) for the TTV PAM sites, 4) performed a least-square fit of the predicted vs. measured growth rate values, and 5) updated the value of the energy matrix only if the least-square fit was smaller than in the previous iteration. The optimized matrix is shown in Fig. 3G.

A similar procedure was followed to extract an optimized energy matrix for the combined TTTV and TTTTV PAM data, obtaining

| (T)TTTV PAM site | -6 | -5 | -4 | -3 | -2 | -1 |
| --- | --- | --- | --- | --- | --- | --- |
| $e^{-\beta\epsilon_T}$ | 0.96 | 1.00 | 1.03 | 1.00 | 1.00 | 0.23 |
| $e^{-\beta\epsilon_C}$ | 1.10 | 1.25 | 0.41 | 0.00 | 0.00 | 0.99 |
| $e^{-\beta\epsilon_G}$ | 0.96 | 0.98 | 0.64 | 0.00 | 0.00 | 0.99 |
| $e^{-\beta\epsilon_A}$ | 0.97 | 0.98 | 0.29 | 0.00 | 0.00 | 0.99 |

The predicted values for the TTV PAM sited were computed from the energy matrix shown in Fig. 3G and those of the form (T)TTTV were computed from the *(T)TTTV PAM site* matrix above. The combined model has a predicted-measured Person correlation of 0.943.

We performed the same procedure to generate the mismatch energy matrix in Fig. 4E. In this case, we first computed the baseline position-dependent energy costs 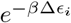 from the values of the growth rate for individual mutations (Fig. 4C). This unoptimized energy matrix is given by

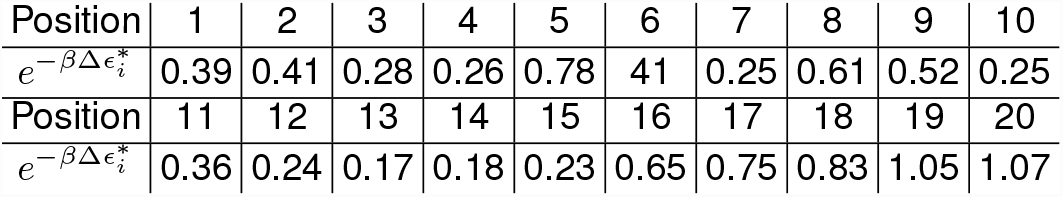

We used this initial set of values for 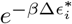 to evaluate the accuracy of the model prediction in computing the growth rate for crRNA that contain two mismatches from 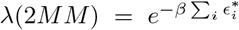. In this case, the Pearson correlation between the predicted and measured growth rate values is 0.769. By subjecting the position-dependent energy matrix to the fitting/optimization procedure described above for 10,000 steps, we obtained the set of values for 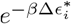 shown in Fig. 4F, which yield a Pearson correlation of 0.869.

---

**FIG. S1.**
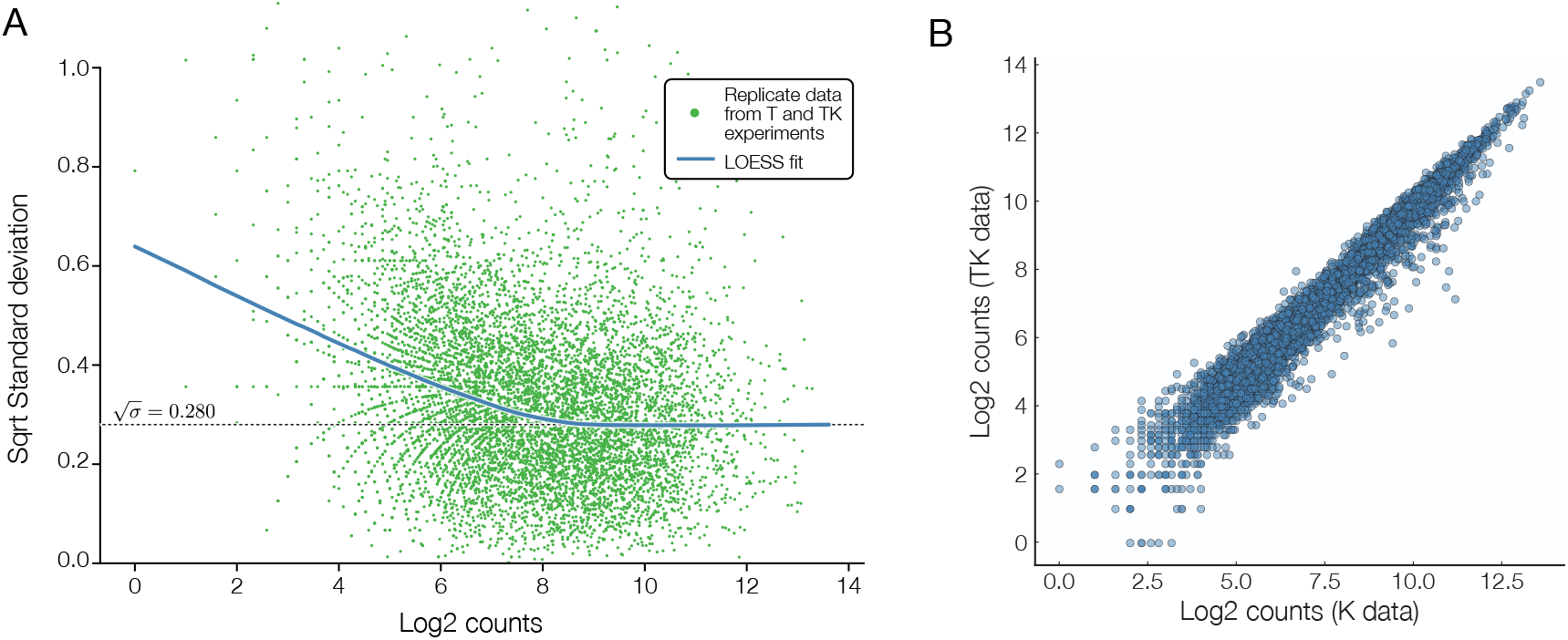
Error calculations. A) Error bars are calculated using a LOESS fit [5] of the mean/variance relationship between experimental replicates of the fold change, inspired by the error estimation in [4]. In this figure, the standard deviation is computed from independent replicates sourced from the K and TK selection conditions with 6 mismatches in the seed sequence. B) Comparing the raw counts from the K and TK independent replicates.

**FIG. S2.**
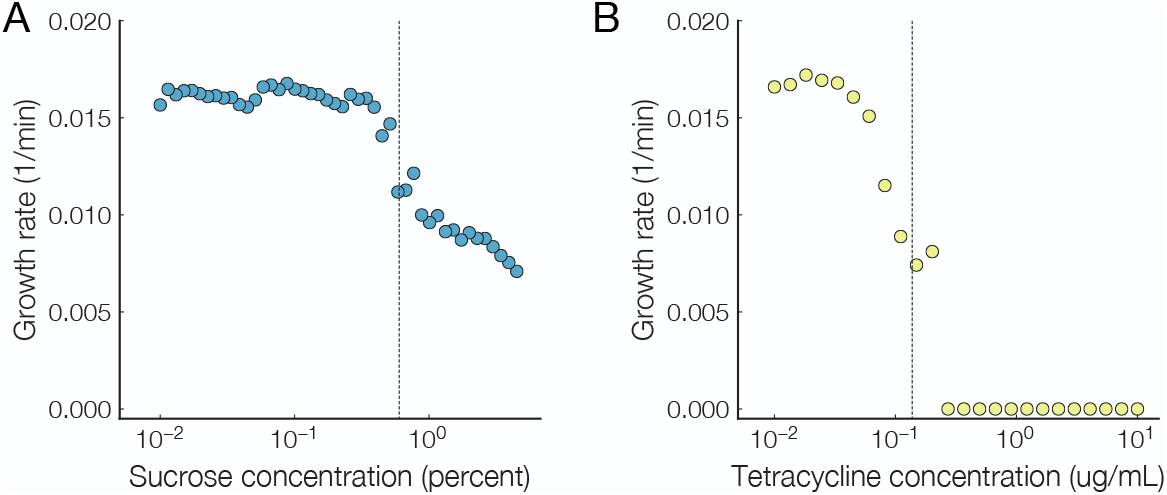
Growth rate under sucrose and tetracycline. A) Sucrose-dependent growth rate for cells that fully express the tetA-sacB cassette. Transition between growth/no-growth occurs at 0.6%. B) Tetracycline-dependent growth rate for cells that fully express the tetA-sacB cassette. Transition between no-growth/growth occurs at 0.14*μ*g/mL.

**FIG. S3.**
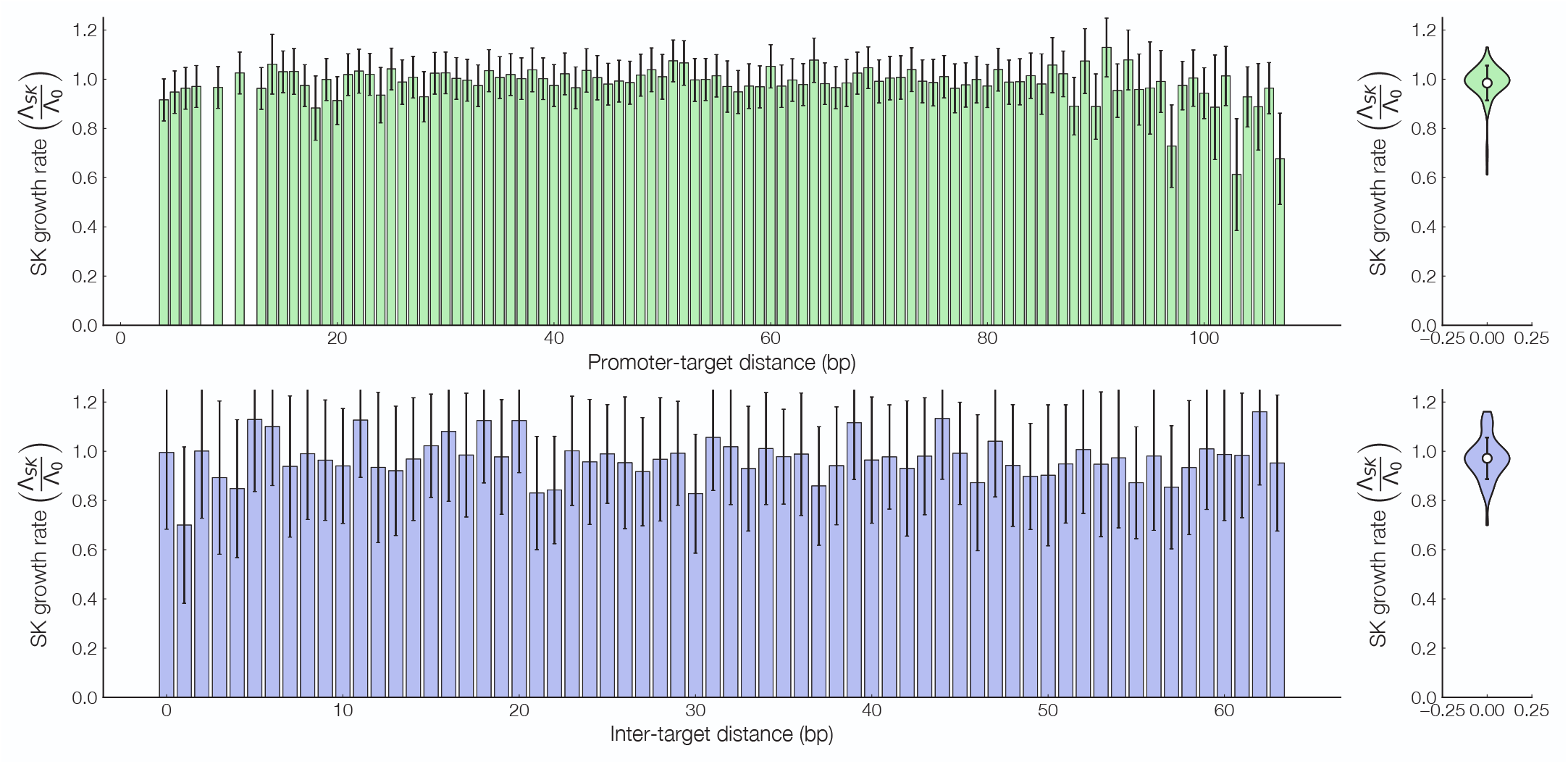
Promoter-target and intra-target distances. Growth rate in SK conditions for inverter constructs which contain (A) a single target or (B) two targets located after the +1 location of the output promoter. Violin plots of the distribution of all constructs is shown on the left. Error bars = std. dev.

**FIG. S4.**
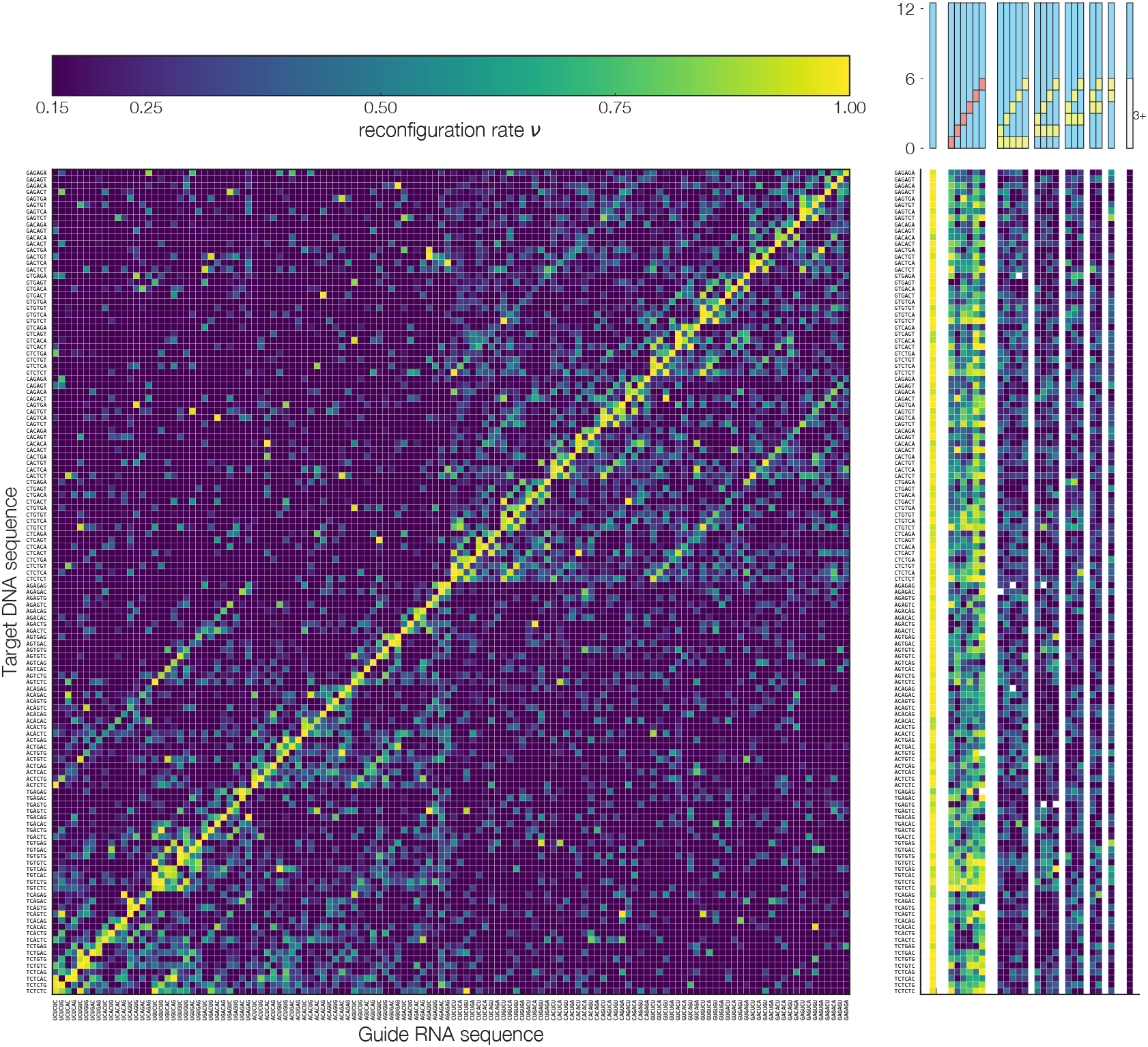
Cross-talk map target DNA dependent mismatch map of the growth rate under 4.5% sucrose (SK) conditions for (SWSWSW+WSWSWS)×(SWSWSW+WSWSWS) seed constructs.

**FIG. S5.**
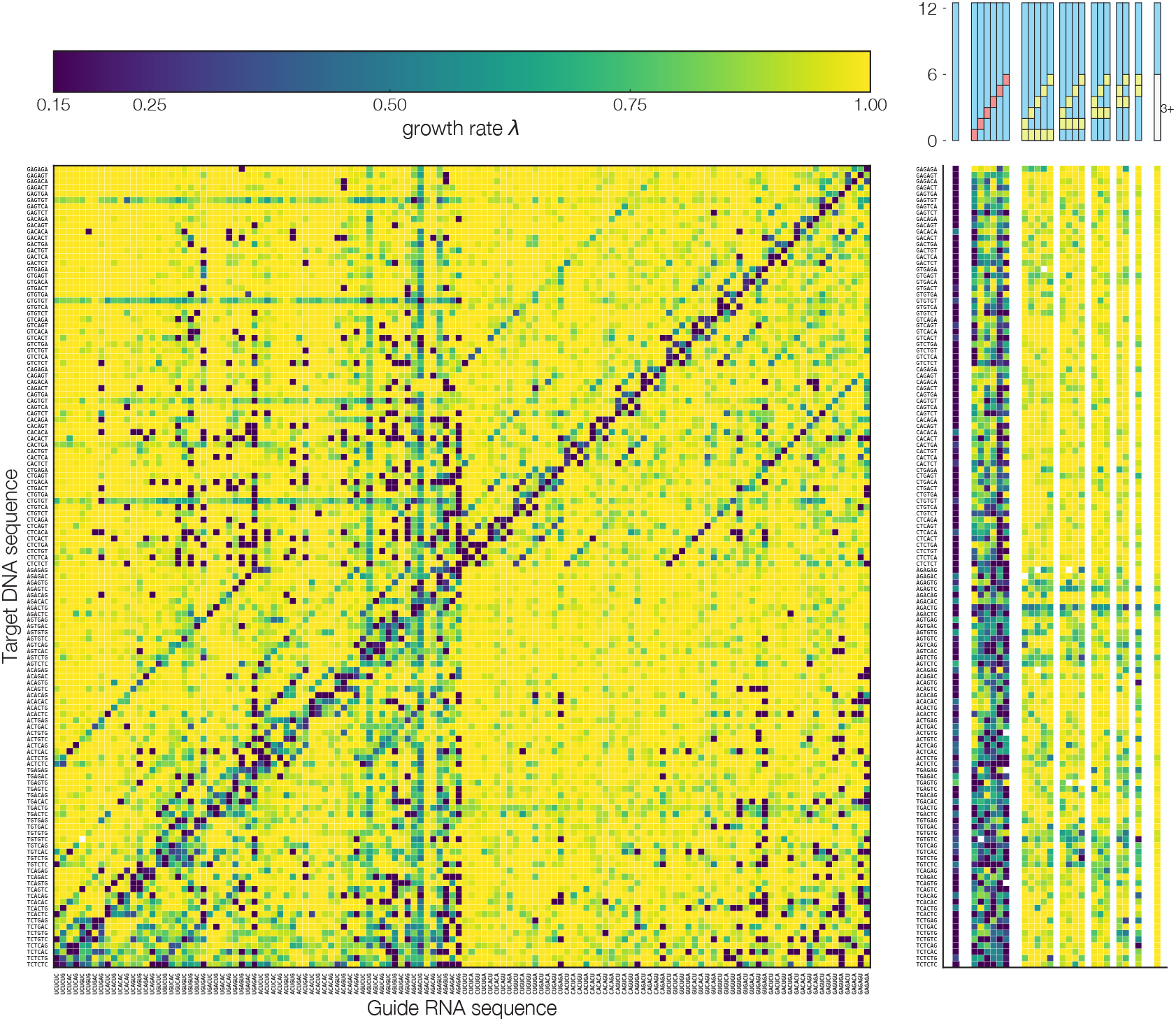
Cross-talk map target DNA dependent mismatch map of the growth rate under 10*μ*g/mL tetracycline (TK) conditions for (SWSWSW+WSWSWS)×(SWSWSW+WSWSWS) seed constructs.

**FIG. S6.**
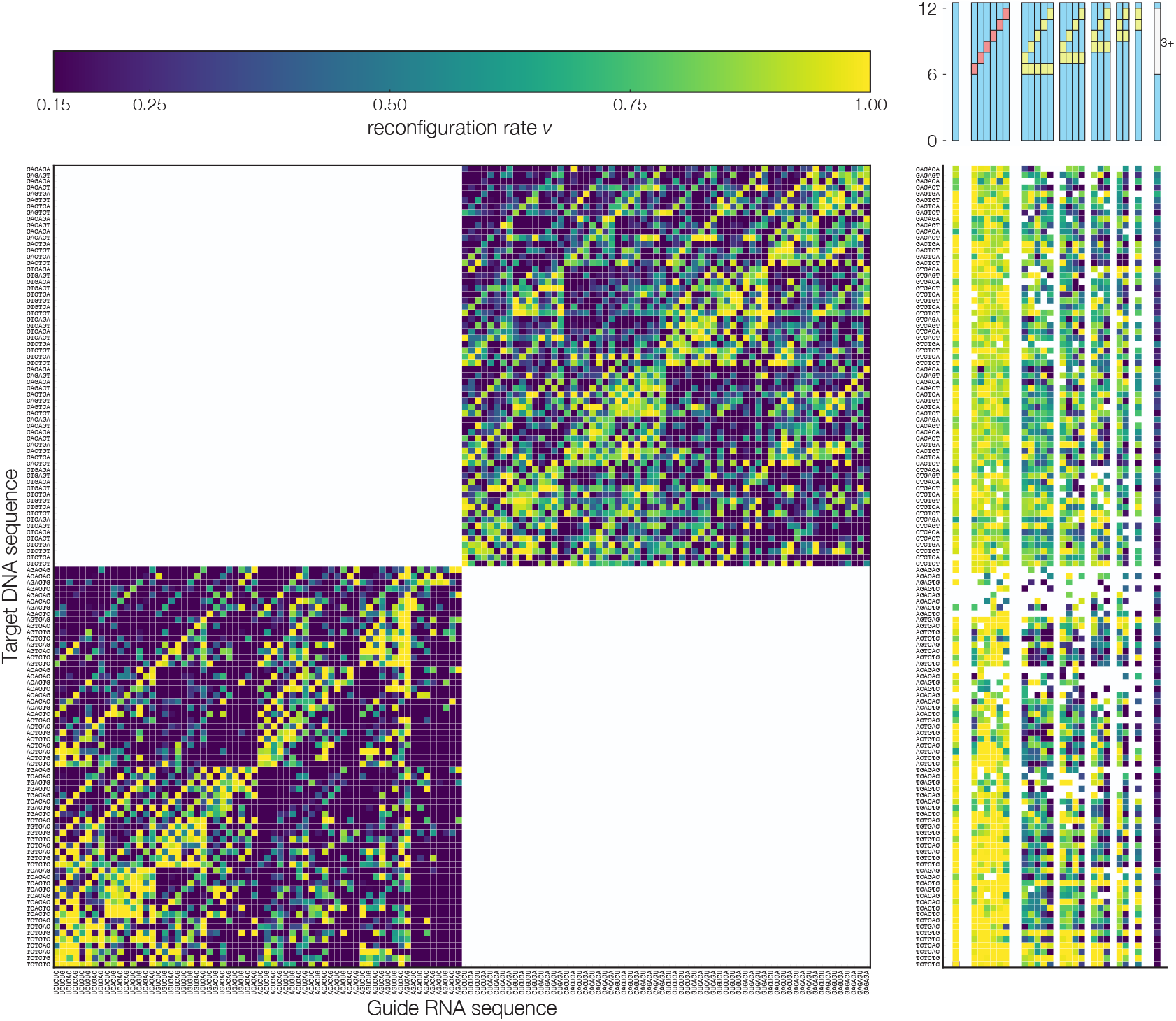
Cross-talk map target DNA dependent mismatch map of the growth rate under 4.5% sucrose (SK) conditions for (SWSWSW)×(SWSWSW) + (WSWSWS)×(WSWSWS) trunk constructs.

**FIG. S7.**
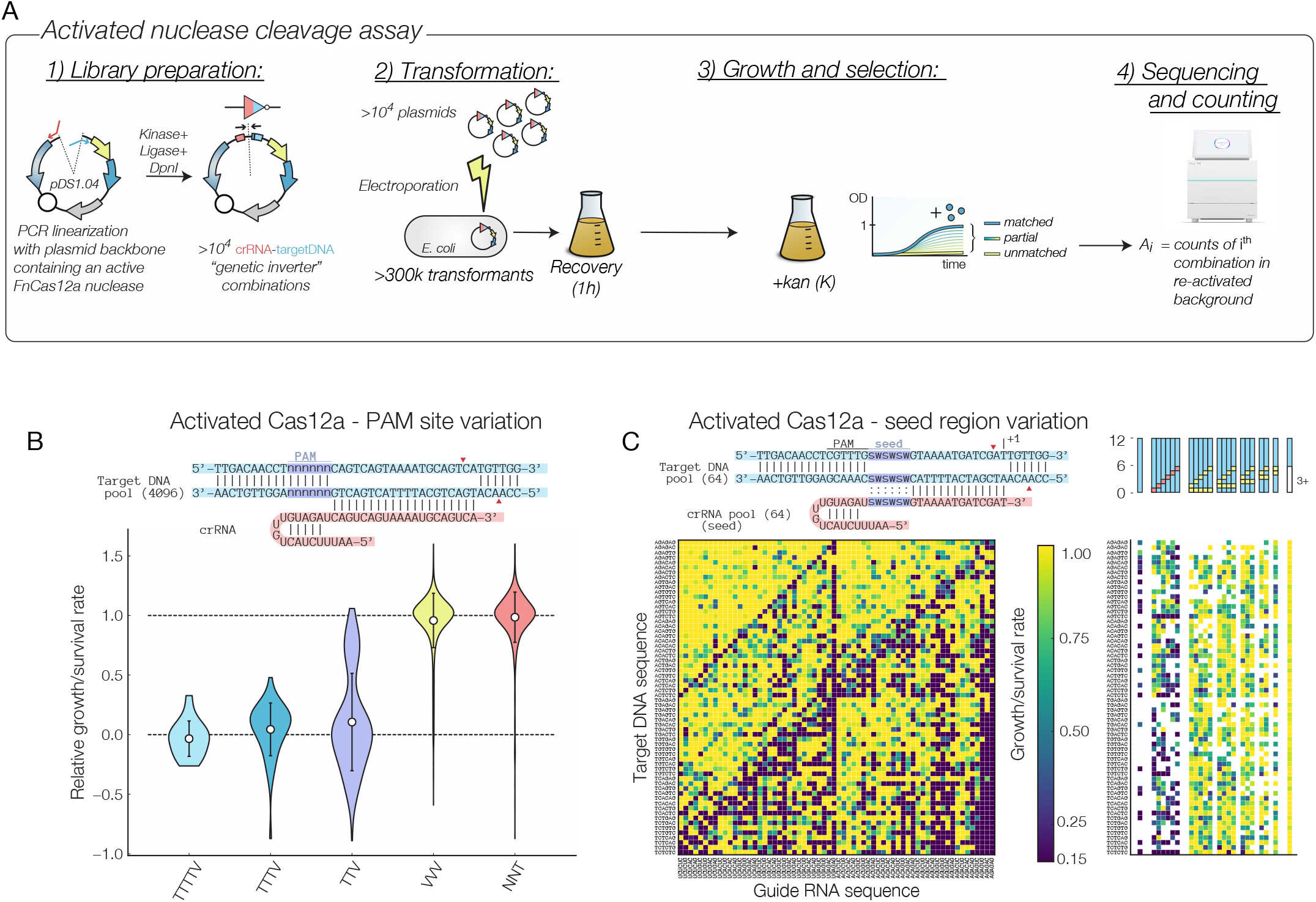
A) Experimental workflow for experiment with activated FnCas12a nuclease sites. B) Relative growth/survival rates for activate FnCas12a nuclease that targets a DNA sequence with a 5’-NNNNNN-3’ PAM site located at the promoter’s −19 position. C) Cross-talk map target DNA dependent mismatch map of the relative density of (SWSWSW+WSWSWS) seed constructs for FnCas12a with re-activated nuclease sites.

## Notes

#### Summary of Updates

Updated Figure 3 and Supplementary Figure 7.

https://www.ncbi.nlm.nih.gov/Traces/study/?acc=PRJNA549693

