## Supplementary material for "Massively parallel CRISPRi assays reveal concealed thermodynamic determinants of dCas12a binding": 12SW-Analysis.ipynb

```
# %% [code]
%pylab
from collections import OrderedDict
import json
import os
from Bio import SeqIO
from scipy.interpolate import interp1d

%matplotlib inline

def reverseComplement(seq):
    complement = {'A': 'T', 'C': 'G', 'G': 'C', 'T': 'A'}
    seq = seq.upper()
    return "".join(complement.get(base, base) for base in reversed(seq))

def Complement(seq):
    complement = {'A': 'T', 'C': 'G', 'G': 'C', 'T': 'A'}
    seq = seq.upper()
    return "".join(complement.get(base, base) for base in (seq))

FILEPATH = '../fastq/'

counts = loadtxt('../error/counts.txt', delimiter= '\n', unpack = False)
logerror = loadtxt('../error/logerror.txt', delimiter = '\n', unpack = False)

#We assume that this then asymtotes to the final value - so we add:
counts = np.append(counts, [1e+07])
logerror = np.append(logerror, [0.07829763])

ferr = interp1d(counts,logerror)

# %% [code]
def importfastq(fname):
    handle1 = open(fname, "rU")
    readiter1 = SeqIO.parse(handle1, "fastq")
    BC1list = [None] * 50*10**6
    k1 = 0
    for r1 in readiter1:
        BC1list[k1] = str(r1.seq)
        k1 += 1
        if (k1 % 1000000)==0:
            print(k1)
    return BC1list[:k1]


files = os.listdir(FILEPATH)

#Finding fastq files
fastq = []
for f in files:
    if f.endswith('R1.fastq') or f.endswith('R1_001.fastq'):
        fastq.append(FILEPATH+f)

#Reading fastq files

for f1 in fastq:
    f2t = f1.split('R1')
    #Check if file exists
    if not os.path.exists(FILEPATH+f2t[0][:-1]+'_DUMB.json'):
        f2 = f2t[0]+'R2'+f2t[1]
        BC1 = importfastq(f1)
        BC2 = importfastq(f2)
        # Add counts to dictionary
        dictionary = {}
        for i in range(len(BC1)):
            guide = BC1[i]+'_'+reverseComplement(BC2[i])
            if guide in dictionary:
                dictionary[guide] += 1
            else:
                dictionary[guide] = 1
        # Order dictionary by counts        
        dictionary = OrderedDict(sorted(dictionary.items(), key=lambda t: -t[1]))
        # Save dictionary
        with open('../fastq/'+f2t[0][:-1]+'_DUMB.json', 'w') as fp:
            json.dump(dictionary, fp)

# %% [code]
MidK = json.load(open('../fastq/9420_9824_73337_BNR3B_8Mid_K_CGATGT_DUMB.json'))
MidSK = json.load(open('../fastq/9420_9824_73338_BNR3B_8Mid_SK_TGACCA_DUMB.json'))

MidK = OrderedDict(sorted(MidK.items(), key=lambda t: -t[1]))
MidSK = OrderedDict(sorted(MidSK.items(), key=lambda t: -t[1]))


SW66K = json.load(open('../fastq/9420_9824_73339_BNR3B_12SW_K_ACAGTG_DUMB.json'))
SW66SK = json.load(open('../fastq/9420_9824_73340_BNR3B_12SW_SK_CTTGTA_DUMB.json'))

SW66K = OrderedDict(sorted(SW66K.items(), key=lambda t: -t[1]))
SW66SK = OrderedDict(sorted(SW66SK.items(), key=lambda t: -t[1]))

D = [MidK, MidSK, SW66K, SW66SK]

# %% [code]
S = ['G', 'C']
W = ['A', 'T']
#SWWWWWWSSWSSWW
dictGuide8 = {}
dictGuide12 = {}
k = 0
for s1 in S:
    for s2 in W:
        for s3 in W:
            for s4 in W:
                for s5 in W:
                    for s6 in W:
                        for s7 in W:
                            for s8 in S:
                                guide = 'CAGTCA'+s1+s2+s3+s4+s5+s6+s7+s8+'CAGTCA'
                                dictGuide8[guide] = k
                                k += 1
k = 0
for s1 in S:
    for s2 in W:
        for s3 in S:
            for s4 in W:
                for s5 in S:
                    for s6 in W:
                        for s7 in S:
                            for s8 in W:
                                for s9 in S:
                                    for s10 in W:
                                        for s11 in S:
                                            for s12 in W:
                                                guide = s1+s2+s3+s4+s5+s6+'GTAAAATG'+s7+s8+s9+s10+s11+s12
                                                dictGuide12[guide] = k
                                                k += 1                                

                        

dictGuide8 = OrderedDict(sorted(dictGuide8.items(), key=lambda t: -t[1])) 
dictGuide12 = OrderedDict(sorted(dictGuide12.items(), key=lambda t: -t[1])) 

# %% [code]
# Cas12-Repression
ν
dicts = []
sumVals = range(4)
for i in range(4):
    dictAll = {}
    for key, value in D[i].items():
        try:
            if (key.find('TGACAGACAATCGACTGCGTAGTAAAATCCCTAAACAATTTCTACTGTTGTAGAT')>-1 and
                key.find('TTTTGGAGCCTTTTTTTTTTGACAACCTCGTTTGCAGTCAGTAAAATGCAGTCATGTTGGGGATGAAGAAATGTAC') > -1):
                bc = key.split('AATTTCTACTGTTGTAGAT')[1][:20]
                if (bc in dictGuide8 and i <= 1) or (bc in dictGuide12 and i >= 2):
                    if bc in dictAll:
                        dictAll[bc] += double(value)
                    else:
                        dictAll[bc] = double(value)
        except:
            pass
    dictAll = OrderedDict(sorted(dictAll.items(), key=lambda t: -t[1]))
    dicts.append(OrderedDict(dictAll))

for i in range(4):
    sumVals[i] = sum(dict(dicts[i]).values())
    for key, value in dicts[i].items():
        dicts[i][key] = value/sumVals[i]


# %% [code]
def extractGrowthRate(dictK, dictSK, fdist=6, idx=(1,0), canon = 'CAGTCAGTAAAATGCAGTCA'):
    RepN = []  # No growth control
    RepF = []  # Full Growth control

    for key, value in dictSK.items():
        hdist = sum(c1 != c2 for c1, c2 in zip(key[:-3], canon[:-3]))
        if hdist >= fdist:
            RepN.append(key)
        elif hdist == 0:
            RepF.append(key)


    ## Find Tk, time cells grew in K-selection
    sm = 0
    km = 0
    tt = 0
    N = 0
    for k in RepN:
        try:
            if dictSK[k] < dictK[k]:
                sm += dictSK[k]
                km += dictK[k]
                N += 1
        except:
            pass 
    Tk = -log(sm/km)

    ## Find Tk, time cells grew in SK-selection
    sp = 0
    kp = 0
    ttt = 0
    N = 0
    for k in RepF:
        try:
            sp += dictSK[k]
            kp += dictK[k]
            N += 1
        except:
            pass
    Ts = log(sp/kp)+Tk

    RepG = {}
    RepGerr = {}      
    for key, value in dictK.items():
            try:
                RepG[key] = (log(dictSK[key]/dictK[key])- log(sm/km))/(log(sp/kp) - log(sm/km))
                RepGerr[key] = sqrt(ferr(dictSK[key]*sumVals[idx[0]])**2+ferr(dictK[key]*sumVals[idx[1]])**2)/log(2)/(log(sp/kp) - log(sm/km))
            except:
                pass
    RepG = OrderedDict(sorted(RepG.items(), key=lambda t: -t[1]))        
    return RepG, RepGerr

##########
## Mid8 Dataset
##########

RepG8, RepG8err = extractGrowthRate(dicts[0], dicts[1], 7, (1,0))
RepG12, RepG12err = extractGrowthRate(dicts[2], dicts[3], 7, (3,2))

RepE = {}
for key, value in RepG12.items():
    valueRescaled = max(0.001, min(value, 0.995))
    RepE[key] = -log(1-valueRescaled)
for key, value in RepG8.items():
    valueRescaled = max(0.001, min(value, 0.995))
    RepE[key] = -log(1-valueRescaled)
    
RepE = OrderedDict(sorted(RepE.items(), key=lambda t: -t[1]))


# %% [code]
Data = []
Data.append(['gRNA-sequence', 'Counts-K', 'Fraction-K', 'Counts-SK', 'Fraction-SK', 'GrowthRate', 'GrowthRateError', 'BindingEnergy'])
for key, value in dicts[0].items():
    try:
        Data.append([key, 
                     str(round(value*sumVals[0])), str(value), 
                     str(round(dicts[1][key]*sumVals[1])), str(dicts[1][key]), 
                     str(RepG8[key]), str(RepG8err[key]), str(RepE[key])])
    except:
        pass

Data.append(['gRNA-sequence', 'Counts-K', 'Fraction-K', 'Counts-SK', 'Fraction-SK', 'GrowthRate', 'GrowthRateError', 'BindingEnergy'])
for key, value in dicts[2].items():
    try:
        Data.append([key, 
                     str(round(value*sumVals[2])), str(value), 
                     str(round(dicts[3][key]*sumVals[1])), str(dicts[3][key]), 
                     str(RepG12[key]), str(RepG12err[key]), str(RepE[key])])
    except:
        pass    
    
f = open("../csv/BinaryMismatchData.csv","w+")
for s in Data:
    f.write(','.join(s)+'\n')
f.close()   

save('../npy/BinaryMismatchData.npy', Data)
```
