## Supplementary material for "Massively parallel CRISPRi assays reveal concealed thermodynamic determinants of dCas12a binding": First6-Analysis.ipynb

```
# %% [code]
%pylab
from collections import OrderedDict
import json
import os
from Bio import SeqIO
from scipy.interpolate import interp1d

%matplotlib inline

def reverseComplement(seq):
    complement = {'A': 'T', 'C': 'G', 'G': 'C', 'T': 'A'}
    seq = seq.upper()
    return "".join(complement.get(base, base) for base in reversed(seq))

def Complement(seq):
    complement = {'A': 'T', 'C': 'G', 'G': 'C', 'T': 'A'}
    seq = seq.upper()
    return "".join(complement.get(base, base) for base in (seq))

FILEPATH = '../fastq/'

counts = loadtxt('../error/counts.txt', delimiter= '\n', unpack = False)
logerror = loadtxt('../error/logerror.txt', delimiter = '\n', unpack = False)

#We assume that this then asymtotes to the final value - so we add:
counts = np.append(counts, [1e+07])
logerror = np.append(logerror, [0.07829763])

ferr = interp1d(counts,logerror)

# %% [code]
def importfastq(fname):
    handle1 = open(fname, "rU")
    readiter1 = SeqIO.parse(handle1, "fastq")
    BC1list = [None] * 50*10**6
    k1 = 0
    for r1 in readiter1:
        BC1list[k1] = str(r1.seq)
        k1 += 1
        if (k1 % 1000000)==0:
            print(k1)
    return BC1list[:k1]


files = os.listdir(FILEPATH)

#Finding fastq files
fastq = []
for f in files:
    if f.endswith('R1.fastq') or f.endswith('R1_001.fastq'):
        fastq.append(FILEPATH+f)

#Reading fastq files

for f1 in fastq:
    f2t = f1.split('R1')
    #Check if file exists
    if not os.path.exists(FILEPATH+f2t[0][:-1]+'_DUMB.json'):
        f2 = f2t[0]+'R2'+f2t[1]
        BC1 = importfastq(f1)
        BC2 = importfastq(f2)
        # Add counts to dictionary
        dictionary = {}
        for i in range(len(BC1)):
            guide = BC1[i]+'_'+reverseComplement(BC2[i])
            if guide in dictionary:
                dictionary[guide] += 1
            else:
                dictionary[guide] = 1
        # Order dictionary by counts        
        dictionary = OrderedDict(sorted(dictionary.items(), key=lambda t: -t[1]))
        # Save dictionary
        with open('../fastq/'+f2t[0][:-1]+'_DUMB.json', 'w') as fp:
            json.dump(dictionary, fp)

# %% [code]
AK = json.load(open('../fastq/9124_9824_67636_BKBLR_cas12_WS_K_ACTTGA_DUMB.json'))
AK = OrderedDict(sorted(AK.items(), key=lambda t: -t[1]))

ASK = json.load(open('../fastq/9124_9824_67638_BKBLR_cas12_WS_SK_TAGCTT_DUMB.json'))
ASK = OrderedDict(sorted(ASK.items(), key=lambda t: -t[1]))

ATK = json.load(open('../fastq/9124_9824_67637_BKBLR_cas12_WS_TK_GATCAG_DUMB.json'))
ATK = OrderedDict(sorted(ATK.items(), key=lambda t: -t[1]))

D1 = [AK, ASK, ATK]

BK = json.load(open('../fastq/9295_9824_71339_BLTF9_K_TGACCA_DUMB.json'))
BK = OrderedDict(sorted(BK.items(), key=lambda t: -t[1]))

BSK = json.load(open('../fastq/9295_9824_71340_BLTF9_SK_ACAGTG_DUMB.json'))
BSK = OrderedDict(sorted(BSK.items(), key=lambda t: -t[1]))

BTK = json.load(open('../fastq/9295_9824_71341_BLTF9_TK_CTTGTA_DUMB.json'))
BTK = OrderedDict(sorted(BTK.items(), key=lambda t: -t[1]))

D2 = [BK, BSK, BTK]

CK = json.load(open('../fastq/9562_9824_74783_BKLHV_cas12_SW_K_CGATGT_DUMB.json'))
CK = OrderedDict(sorted(CK.items(), key=lambda t: -t[1]))

CSK = json.load(open('../fastq/9562_9824_74784_BKLHV_cas12_SW_SK_TGACCA_DUMB.json'))
CSK = OrderedDict(sorted(CSK.items(), key=lambda t: -t[1]))

CTK = json.load(open('../fastq/9562_9824_74785_BKLHV_cas12_SW_TK_ACAGTG_DUMB.json'))
CTK = OrderedDict(sorted(CTK.items(), key=lambda t: -t[1]))

D3 = [CK, CSK, CTK]

# %% [code]
D = [AK, ASK, ATK,
     BK, BSK, BTK,
     CK, CSK, CTK]

# %% [code]
S = ['G', 'C']
W = ['A', 'T']
#SWWWWWWSSWSSWW
dictGuide = {}
dictGuideSW = {}
dictGuideWS = {}
k = 0
for s1 in S:
    for s2 in W:
        for s3 in S:
            for s4 in W:
                for s5 in S:
                    for s6 in W:
                        guide = s1+s2+s3+s4+s5+s6
                        dictGuideSW[guide] = k
                        dictGuide[guide] = k
                        k += 1
for s1 in W:
    for s2 in S:
        for s3 in W:
            for s4 in S:
                for s5 in W:
                    for s6 in S:
                        guide = s1+s2+s3+s4+s5+s6
                        dictGuideWS[guide] = k
                        dictGuide[guide] = k
                        k += 1
                        

dictGuideSW = OrderedDict(sorted(dictGuideSW.items(), key=lambda t: -t[1])) 
dictGuideWS = OrderedDict(sorted(dictGuideWS.items(), key=lambda t: -t[1]))
dictGuide = OrderedDict(sorted(dictGuide.items(), key=lambda t: -t[1]))

# %% [code]
# Cas12-Repression

dicts = []
sumVals = range(9)
for i in range(9):
    dictAll = {}
    for key, value in D[i].items():
        if len(key) > 5:
            if (key.find('TGACAGACAATCGACTGCGTAGTAAAATCCCTAAACAATTTCTACTGTTGTAGAT')>-1 and
                key.find('GTAAAATGATCGATA_TTTTGGAGCCTTTTTTTTTTGACAACCTCGTTTG') > -1 and
                key.find('GTAAAATGATCGATTGTTGGGGATGAAGAAATGTAC') > -1):
                bc1 = key.split('AATTTCTACTGTTGTAGAT')[1][:20]
                bc2 = key.split('TTTTGGAGCCTTTTTTTTTTGACAACCTCGTTTG')[1][:20]
                if bc1[:6] in dictGuide and bc2[:6] in dictGuide:
                    if bc1+'_'+bc2 in dictAll:
                        dictAll[bc1+'_'+bc2] += double(value)
                    else:
                        dictAll[bc1+'_'+bc2] = double(value)

    dictAll = OrderedDict(sorted(dictAll.items(), key=lambda t: -t[1]))
    dicts.append(OrderedDict(dictAll))

    

for i in range(9):
    sumVals[i] = sum(dict(dicts[i]).values())
    for key, value in dicts[i].items():
        dicts[i][key] = value/sumVals[i]


# %% [code]
Xall = range(9)

for i in range(9):
    Xall[i] = zeros((128, 128))
    for key, value in dicts[i].items():
        bc1 = key.split('GTAAAATGATCGAT_')[0]
        bc2 = key.split('GTAAAATGATCGAT_')[1][:6]
        Xall[i][dictGuide[bc1], dictGuide[bc2]] += value

# %% [code]
def extractGrowthRate(dictK, dictSK, hselect, idx=(1,0)):
    RepN = []  # No growth control
    RepF = []  # Full Growth control
    canon = 'CAGTCAGTAAAATGCAGTCA'

    for key, value in dictSK.items():
        try:
            bc1, bc2 = key.split('_')
            hdist = sum(c1 != c2 for c1, c2 in zip(bc1, bc2))
            if hdist == hselect[0]:
                RepN.append(key)
            elif hdist == hselect[1]:
                RepF.append(key)
        except:
            pass  

    ## Find Tk, time cells grew in K-selection
    sm = 0
    km = 0
    tt = 0
    N = 0
    for k in RepN:
        try:
            sm += dictSK[k]
            km += dictK[k]
            N += 1
        except:
            pass
    Tk = -log(sm/km)

    ## Find Tk, time cells grew in SK-selection
    sp = 0
    kp = 0
    ttt = 0
    N = 0
    for k in RepF:
        try:
            sp += dictSK[k]
            kp += dictK[k]
            N += 1
        except:
            pass
    Ts = log(sp/kp)+Tk

    RepG = {}
    RepGerr = {}    
    for key, value in dictK.items():
            try:
                RepG[key] = (log(dictSK[key]/dictK[key])- log(sm/km))/(log(sp/kp) - log(sm/km))
                RepGerr[key] = sqrt(ferr(dictSK[key]*sumVals[idx[1]])**2+ferr(dictK[key]*sumVals[idx[0]])**2)/log(2)/(log(sp/kp) - log(sm/km))                                
            except:
                pass
    RepG = OrderedDict(sorted(RepG.items(), key=lambda t: -t[1]))        
    return RepG, RepGerr

##########
## Mid8 Dataset
##########

RepGWS, RepGWSerr = extractGrowthRate(dicts[3], dicts[4], [6, 0], (3, 4))
RepGSW, RepGSWerr = extractGrowthRate(dicts[6], dicts[7], [6, 0], (6, 7))


RepE = {}
for key, value in RepGWS.items():
    bc1, bc2 = key.split('_')
    if bc1[:6] in dictGuideWS:    
        valueRescaled = max(0.001, min(value, 0.995))
        RepE[key] = -log(1-valueRescaled)
for key, value in RepGSW.items():
    bc1, bc2 = key.split('_')
    if bc1[:6] in dictGuideSW:    
        valueRescaled = max(0.001, min(value, 0.995))
        RepE[key] = -log(1-valueRescaled)

    
RepES = OrderedDict(sorted(RepE.items(), key=lambda t: -t[1]))


RepGWST,RepGWSTerr  = extractGrowthRate(dicts[3], dicts[5], [0, 6], [3,5])
RepGSWT,RepGSWTerr = extractGrowthRate(dicts[6], dicts[8], [0, 6], [3,5])

'''
RepE = {}
for key, value in RepGWST.items():
    bc1, bc2 = key.split('_')
    if bc1[:6] in dictGuideWS:
        valueRescaled = max(0.001, min(value, 0.995))
        RepE[key] = -log(1-valueRescaled)
for key, value in RepGSWT.items():
    bc1, bc2 = key.split('_')
    if bc1[:6] in dictGuideSW:
        valueRescaled = max(0.001, min(value, 0.995))
        RepE[key] = -log(1-valueRescaled)


RepET = OrderedDict(sorted(RepE.items(), key=lambda t: -t[1]))
'''

# %% [code]
Data = []
Data.append(['Barcode', 'Counts-K', 'Fraction-K', 'Counts-SK', 'Fraction-SK', 'Counts-TK', 'Fraction-TK', 'GrowthRate-SK', 'GrowthRate-SK-Error', 'Energy-SK', 'GrowthRate-TK', 'GrowthRate-TK-Error'])
for key, value in dicts[3].items():
    bc1, bc2 = key.split('_')
    try:
        if bc1[:6] in dictGuideWS:
            Data.append([key, 
                         str(round(value*sumVals[3])), str(value), 
                         str(round(dicts[4][key]*sumVals[4])), str(dicts[4][key]), 
                         str(round(dicts[5][key]*sumVals[5])), str(dicts[5][key]), 
                         str(RepGWS[key]), str(RepGWSerr[key]), str(RepES[key]),
                         str(RepGWST[key]), str(RepGWSTerr[key])])
    except:
        pass
for key, value in dicts[6].items():
    bc1, bc2 = key.split('_')
    try:
        if bc1[:6] in dictGuideSW:
            Data.append([key, 
                         str(round(value*sumVals[6])), str(value), 
                         str(round(dicts[7][key]*sumVals[7])), str(dicts[7][key]), 
                         str(round(dicts[8][key]*sumVals[8])), str(dicts[8][key]), 
                         str(RepGSW[key]), str(RepGSWerr[key]), str(RepES[key]),
                         str(RepGSWT[key]), str(RepGSWTerr[key])])
    except:
        pass

f = open("First6-Data.csv","w+")
for s in Data:
    f.write(','.join(s)+'\n')
f.close()   

save("First6-Data.npy", Data)

# %% [code]
Data2 = []
Data2.append(['Barcode', 'Counts-K', 'Fraction-K', 'Counts-SK', 'Fraction-SK', 'Counts-TK', 'Fraction-TK', 'Fitness'])
for key, value in dicts[3].items():
    try:
        
        Data2.append([key, 
                     str(round(value*sumVals[0])), str(value), 
                     str(round(dicts[1][key]*sumVals[1])), str(dicts[1][key]), 
                     str(round(dicts[2][key]*sumVals[2])), str(dicts[2][key]), 
                     str(log(dicts[3][key]/dicts[2][key])/log(2))])
    except:
        pass
for key, value in dicts[0].items():
    try:
        Data2.append([key, 
                     str(round(value*sumVals[0])), str(value), 
                     str(round(dicts[1][key]*sumVals[1])), str(dicts[1][key]), 
                     str(round(dicts[2][key]*sumVals[2])), str(dicts[2][key]), 
                     str(log(dicts[7][key]/dicts[8][key])/log(2))])
    except:
        pass

f = open("../csv/First6-ExtraDuplicateControl-Data.csv","w+")
for s in Data2:
    f.write(','.join(s)+'\n')
f.close()   
save("../npy/First6-ExtraDuplicateControl-Data.npy", Data2)
```
