## Supplementary material for "Massively parallel CRISPRi assays reveal concealed thermodynamic determinants of dCas12a binding": Mid6-Analysis.ipynb

```
# %% [code]
%pylab
from collections import OrderedDict
import json
import os
from Bio import SeqIO
from scipy.interpolate import interp1d

%matplotlib inline

def reverseComplement(seq):
    complement = {'A': 'T', 'C': 'G', 'G': 'C', 'T': 'A'}
    seq = seq.upper()
    return "".join(complement.get(base, base) for base in reversed(seq))

def Complement(seq):
    complement = {'A': 'T', 'C': 'G', 'G': 'C', 'T': 'A'}
    seq = seq.upper()
    return "".join(complement.get(base, base) for base in (seq))

FILEPATH = '../fastq'

counts = loadtxt('../error/counts.txt', delimiter= '\n', unpack = False)
logerror = loadtxt('../error/logerror.txt', delimiter = '\n', unpack = False)

#We assume that this then asymtotes to the final value - so we add:
counts = np.append(counts, [1e+07])
logerror = np.append(logerror, [0.07829763])

ferr = interp1d(counts,logerror)

# %% [code]
def importfastq(fname):
    handle1 = open(fname, "rU")
    readiter1 = SeqIO.parse(handle1, "fastq")
    BC1list = [None] * 50*10**6
    k1 = 0
    for r1 in readiter1:
        BC1list[k1] = str(r1.seq)
        k1 += 1
        if (k1 % 1000000)==0:
            print(k1)
    return BC1list[:k1]


files = os.listdir(FILEPATH)

#Finding fastq files
fastq = []
for f in files:
    if f.endswith('R1.fastq') or f.endswith('R1_001.fastq'):
        fastq.append(FILEPATH+f)

#Reading fastq files

for f1 in fastq:
    f2t = f1.split('R1')
    #Check if file exists
    if not os.path.exists(FILEPATH+f2t[0][:-1]+'_DUMB.json'):
        f2 = f2t[0]+'R2'+f2t[1]
        BC1 = importfastq(f1)
        BC2 = importfastq(f2)
        # Add counts to dictionary
        dictionary = {}
        for i in range(len(BC1)):
            guide = BC1[i]+'_'+reverseComplement(BC2[i])
            if guide in dictionary:
                dictionary[guide] += 1
            else:
                dictionary[guide] = 1
        # Order dictionary by counts        
        dictionary = OrderedDict(sorted(dictionary.items(), key=lambda t: -t[1]))
        # Save dictionary
        with open('../fastq/'+f2t[0][:-1]+'_DUMB.json', 'w') as fp:
            json.dump(dictionary, fp)

# %% [code]
WK = json.load(open('../fastq/9793_10347_78396_C25R3_WS_12mid_Kan_CGATGT_DUMB.json'))
WK = OrderedDict(sorted(WK.items(), key=lambda t: -t[1]))

WSK = json.load(open('../fastq/9793_10347_78397_C25R3_WS_12mid_Sucrose_Kan_TGACCA_DUMB.json'))
WSK = OrderedDict(sorted(WSK.items(), key=lambda t: -t[1]))

WTK = json.load(open('../fastq/9793_10347_78398_C25R3_WS_12mid_TetA_Kan_ACAGTG_DUMB.json'))
WTK = OrderedDict(sorted(WTK.items(), key=lambda t: -t[1]))

SK = json.load(open('../fastq/9793_10347_78399_C25R3_SW_12mid_Kan_GCCAAT_DUMB.json'))
SK = OrderedDict(sorted(SK.items(), key=lambda t: -t[1]))

SSK = json.load(open('../fastq/9793_10347_78400_C25R3_SW_12mid_Sucrose_Kan_CAGATC_DUMB.json'))
SSK = OrderedDict(sorted(SSK.items(), key=lambda t: -t[1]))

STK = json.load(open('../fastq/9793_10347_78401_C25R3_SW_12mid_TetA_Kan_CTTGTA_DUMB.json'))
STK = OrderedDict(sorted(STK.items(), key=lambda t: -t[1]))

D = [WK, WSK, WTK,
     SK, SSK, STK]

# %% [code]
S = ['G', 'C']
W = ['A', 'T']
#SWWWWWWSSWSSWW
dictGuide = {}
dictGuideSW = {}
dictGuideWS = {}
k = 0
for s1 in S:
    for s2 in W:
        for s3 in S:
            for s4 in W:
                for s5 in S:
                    for s6 in W:
                        guide = s1+s2+s3+s4+s5+s6
                        dictGuide[guide] = k
                        dictGuideSW[guide] = k
                        k += 1
for s1 in W:
    for s2 in S:
        for s3 in W:
            for s4 in S:
                for s5 in W:
                    for s6 in S:
                        guide = s1+s2+s3+s4+s5+s6
                        dictGuide[guide] = k
                        dictGuideWS[guide] = k
                        k += 1
                        

dictGuide = OrderedDict(sorted(dictGuide.items(), key=lambda t: -t[1])) 

# %% [code]
# Cas12-Repression

dicts = []
sumVals = range(6)
for i in range(6):
    dictAll = {}
    for key, value in D[i].items():
        if len(key) > 5:
            if (key.find('TGACAGACAATCGACTGCGTAGTAAAATCCCTAAACAATTTCTACTGTTGTAGATACACTG')>-1 and
                key.find('GCAGTCAGA_TTTTGGAGCCTTTTTTTTTTGACACTG') > -1 and
                key.find('GCAGTCAGTAAAATGCACGAATGTTGGGGATGAAGAAATGTAC') > -1):
                bc1 = key.split('AATTTCTACTGTTGTAGAT')[1][:20]
                bc2 = key.split('TAAAATGCACGAATGTTGGGGATGAAG')[0][-20:]
                if bc1[6:12] in dictGuide and bc2[6:12] in dictGuide:
                    if bc1+'_'+bc2 in dictAll:
                        dictAll[bc1+'_'+bc2] += double(value)
                    else:
                        dictAll[bc1+'_'+bc2] = double(value)

    dictAll = OrderedDict(sorted(dictAll.items(), key=lambda t: -t[1]))
    dicts.append(OrderedDict(dictAll))

    

for i in range(6):
    sumVals[i] = sum(dict(dicts[i]).values())
    for key, value in dicts[i].items():
        dicts[i][key] = value/sumVals[i]


# %% [code]
Xall = range(9)

for i in range(6):
    Xall[i] = zeros((128, 128))
    for key, value in dicts[i].items():
        bc1 = key.split('GCAGTCAG_ACACTG')[0][-6:]
        bc2 = key.split('GCAGTCAG_ACACTG')[1][:6]
        Xall[i][dictGuide[bc1], dictGuide[bc2]] += value
        
        

# %% [code]
def extractGrowthRate(dictK, dictSK, hselect, idx=(1,0)):
    RepN = []  # No growth control
    RepF = []  # Full Growth control


    for key, value in dictSK.items():
        try:
            bc1, bc2 = key.split('_')
            hdist = sum(c1 != c2 for c1, c2 in zip(bc1, bc2))
            if hdist == hselect[0]:
                RepN.append(key)
            elif hdist == hselect[1]:
                RepF.append(key)
        except:
            pass  

    ## Find Tk, time cells grew in K-selection
    sm = 0
    km = 0
    tt = 0
    N = 0
    for k in RepN:
        try:
            sm += dictSK[k]
            km += dictK[k]
            N += 1
        except:
            pass
    Tk = -log(sm/km)

    ## Find Tk, time cells grew in SK-selection
    sp = 0
    kp = 0
    ttt = 0
    N = 0
    for k in RepF:
        try:
            sp += dictSK[k]
            kp += dictK[k]
            N += 1
        except:
            pass
    Ts = log(sp/kp)+Tk

    RepG = {}
    RepGerr = {}      
    for key, value in dictK.items():
            try:
                RepG[key] = (log(dictSK[key]/dictK[key])- log(sm/km))/(log(sp/kp) - log(sm/km))
                RepGerr[key] = sqrt(ferr(dictSK[key]*sumVals[idx[1]])**2+ferr(dictK[key]*sumVals[idx[0]])**2)/log(2)/(log(sp/kp) - log(sm/km))                                                
            except:
                pass
    RepG = OrderedDict(sorted(RepG.items(), key=lambda t: -t[1]))        
    return RepG, RepGerr  

##########
## Mid8 Dataset
##########

RepGWS,RepGWSerr = extractGrowthRate(dicts[0], dicts[1], [6, 0], [0,1])
RepGSW,RepGSWerr = extractGrowthRate(dicts[3], dicts[4], [6, 0],[3,4])


RepE = {}
for key, value in RepGWS.items():
    bc1, bc2 = key.split('_')
    if bc1[6:12] in dictGuideWS:    
        valueRescaled = max(0.001, min(value, 0.995))
        RepE[key] = -log(1-valueRescaled)
for key, value in RepGSW.items():
    bc1, bc2 = key.split('_')
    if bc1[6:12] in dictGuideSW:    
        valueRescaled = max(0.001, min(value, 0.995))
        RepE[key] = -log(1-valueRescaled)

    
RepES = OrderedDict(sorted(RepE.items(), key=lambda t: -t[1]))


RepGWST,RepGWSTerr = extractGrowthRate(dicts[0], dicts[2], [0, 6])
RepGSWT,RepGSWTerr = extractGrowthRate(dicts[3], dicts[5], [0, 6])

'''
RepE = {}
for key, value in RepGWST.items():
    bc1, bc2 = key.split('_')
    if bc1[:6] in dictGuideWS:
        valueRescaled = max(0.001, min(value, 0.995))
        RepE[key] = -log(1-valueRescaled)
for key, value in RepGSWT.items():
    bc1, bc2 = key.split('_')
    if bc1[:6] in dictGuideSW:
        valueRescaled = max(0.001, min(value, 0.995))
        RepE[key] = -log(1-valueRescaled)


RepET = OrderedDict(sorted(RepE.items(), key=lambda t: -t[1]))
'''

# %% [code]
Data = []
Data.append(['Barcode', 'Counts-K', 'Fraction-K', 'Counts-SK', 'Fraction-SK', 'Counts-TK', 'Fraction-TK', 'GrowthRate-SK', 'GrowthRate-SK-Error', 'Energy-SK', 'GrowthRate-TK','GrowthRate-TK-Error'])
for key, value in dicts[0].items():
    bc1, bc2 = key.split('_')
    try:
        if bc1[6:12] in dictGuideWS:
            Data.append([key, 
                         str(round(value*sumVals[0])), str(value), 
                         str(round(dicts[1][key]*sumVals[1])), str(dicts[1][key]), 
                         str(round(dicts[2][key]*sumVals[2])), str(dicts[2][key]), 
                         str(RepGWS[key]), str(RepGWSerr[key]), str(RepES[key]),
                         str(RepGWST[key]),str(RepGWSTerr[key])])
    except:
        pass
for key, value in dicts[3].items():
    bc1, bc2 = key.split('_')
    try:
        if bc1[6:12] in dictGuideSW:
            Data.append([key, 
                         str(round(value*sumVals[3])), str(value), 
                         str(round(dicts[4][key]*sumVals[4])), str(dicts[4][key]), 
                         str(round(dicts[5][key]*sumVals[5])), str(dicts[5][key]), 
                         str(RepGSW[key]), str(RepGSWerr[key]), str(RepES[key]),
                         str(RepGSWT[key]),str(RepGSWTerr[key])])
    except:
        pass
    
f = open("../csv/Mid6-Data.csv","w+")
for s in Data:
    f.write(','.join(s)+'\n')
f.close()  

save('../npy/Mid6-Data.npy', Data)
```
