## Supplementary material for "Massively parallel CRISPRi assays reveal concealed thermodynamic determinants of dCas12a binding": MigratingTarget-Analysis.ipynb

```
# %% [code]
%pylab
from collections import OrderedDict
import json
import os
from Bio import SeqIO
from scipy.interpolate import interp1d

%matplotlib inline

def reverseComplement(seq):
    complement = {'A': 'T', 'C': 'G', 'G': 'C', 'T': 'A'}
    seq = seq.upper()
    return "".join(complement.get(base, base) for base in reversed(seq))

def Complement(seq):
    complement = {'A': 'T', 'C': 'G', 'G': 'C', 'T': 'A'}
    seq = seq.upper()
    return "".join(complement.get(base, base) for base in (seq))

FILEPATH = '../fastq/'

counts = loadtxt('../error/counts.txt', delimiter= '\n', unpack = False)
logerror = loadtxt('../error/logerror.txt', delimiter = '\n', unpack = False)

#We assume that this then asymtotes to the final value - so we add:
counts = np.append(counts, [1e+07])
logerror = np.append(logerror, [0.07829763])

ferr = interp1d(counts,logerror)

# %% [code]
def importfastq(fname):
    handle1 = open(fname, "rU")
    readiter1 = SeqIO.parse(handle1, "fastq")
    BC1list = [None] * 50*10**6
    k1 = 0
    for r1 in readiter1:
        BC1list[k1] = str(r1.seq)
        k1 += 1
        if (k1 % 1000000)==0:
            print(k1)
    return BC1list[:k1]


files = os.listdir(FILEPATH)

#Finding fastq files
fastq = []
for f in files:
    if f.endswith('R1.fastq') or f.endswith('R1_001.fastq'):
        fastq.append(FILEPATH+f)

#Reading fastq files

for f1 in fastq:
    f2t = f1.split('R1')
    #Check if file exists
    if not os.path.exists(FILEPATH+f2t[0][:-1]+'_DUMB.json'):
        f2 = f2t[0]+'R2'+f2t[1]
        BC1 = importfastq(f1)
        BC2 = importfastq(f2)
        # Add counts to dictionary
        dictionary = {}
        for i in range(len(BC1)):
            guide = BC1[i]+'_'+reverseComplement(BC2[i])
            if guide in dictionary:
                dictionary[guide] += 1
            else:
                dictionary[guide] = 1
        # Order dictionary by counts        
        dictionary = OrderedDict(sorted(dictionary.items(), key=lambda t: -t[1]))
        # Save dictionary
        with open('../fastq/'+f2t[0][:-1]+'_DUMB.json', 'w') as fp:
            json.dump(dictionary, fp)

# %% [code]
Cas12 = json.load(open('../fastq/9681_9824_75902_BNNB6_TargetVarsK_CGATGT_DUMB.json'))
Cas12 = OrderedDict(sorted(Cas12.items(), key=lambda t: -t[1]))
Cas12SK = json.load(open('../fastq/9681_9824_75903_BNNB6_TargetVarsSK_TGACCA_DUMB.json'))
Cas12SK = OrderedDict(sorted(Cas12SK.items(), key=lambda t: -t[1]))
Cas12TK = json.load(open('../fastq/9681_9824_75904_BNNB6_TargetVarsTK_ACAGTG_DUMB.json'))
Cas12TK = OrderedDict(sorted(Cas12TK.items(), key=lambda t: -t[1]))
D = [Cas12, Cas12SK, Cas12TK]

gRNAKeys = {'f_122':'acacaACCTGATCTGCGTCGT'.upper(), 
            'f_123': 'acatcaCCTGATCTGCGTCGT'.upper(),
            'f_130': 'cagtcaTGCGTCGTAAAATGC'.upper(),
            'f_131': 'cagtcaGCGTCGTAAAATGCA'.upper(),
            'f_132': 'cagtcaCGTCGTAAAATGCAC'.upper(),
            'f_133': 'cagtcaGTCGTAAAATGCACG'.upper(),
            'f_134': 'cagtcaTCGTAAAATGCACGA'.upper(),
            'f_135': 'cagtcaCGTAAAATGCACGAA'.upper(),
            'f_136': 'cagtcaGTAAAATGCACGAAC'.upper(),
            'f_137': 'cagtcaTAAAATGCACGAACG'.upper(),
            'f_138': 'cagtctAAAATGCACGAACGT'.upper(),
            'f_139': 'cagttaAAATGCACGAACGTT'.upper(),
            'f_140': 'cagtaaAATGCACGAACGTTG'.upper(),
            'f_141': 'cataaaATGCACGAACGTTGT'.upper(),
            'f_142': 'ctaaaaTGCACGAACGTTGTT'.upper(),
            'f_143': 'taaaatGCACGAACGTTGTTG'.upper(),
            'f_152': 'cagtcaTTGTTGGGGATGAAG'.upper(),
            'r_136': 'CAGTCAAGGTGTTGTCAAAAA'.upper(),
            'r_137': 'CAGTCACAGGTGTTGTCAAAA'.upper(),
            'r_147': 'ATTTTACGACGCAGATCAGGT'.upper(),
            'r_148': 'CATTTTACGACGCAGATCAGG'.upper(),
            'r_149': 'CAATTTTACGACGCAGATCAG'.upper(),
            'r_150': 'CAGATTTTACGACGCAGATCA'.upper(),
            'r_151': 'CAGTATTTTACGACGCAGATC'.upper(),
            'r_152': 'CAGTCATTTTACGACGCAGAT'.upper(),
            'r_153': 'CAGTCAATTTTACGACGCAGA'.upper()}

for key, value in dict(gRNAKeys).items():
    gRNAKeys[value] = key

tDNAKeys = {'f_122': 'TTTTGACACAACCTGATCTGCGTCGTAAAATGCACGAACGT',
            'f_123': 'TTTTGACATCACCTGATCTGCGTCGTAAAATGCACGAACGT',
            'f_130': 'TTTTGACATTTGCAGTCATGCGTCGTAAAATGCACGAACGT',
            'f_131': 'TTTTGACAATTTGCAGTCAGCGTCGTAAAATGCACGAACGT',
            'f_132': 'TTTTGACAACTTTGCAGTCACGTCGTAAAATGCACGAACGT',
            'f_133': 'TTTTGACAACATTTGCAGTCAGTCGTAAAATGCACGAACGT',
            'f_134': 'TTTTGACAACACTTTGCAGTCATCGTAAAATGCACGAACGT',
            'f_135': 'TTTTGACAACACCTTTGCAGTCACGTAAAATGCACGAACGT',
            'f_136': 'TTTTGACAACACCTTTTGCAGTCAGTAAAATGCACGAACGT',
            'f_137': 'TTTTGACAACACCTGTTTGCAGTCATAAAATGCACGAACGT',
            'f_138': 'TTTTGACAACACCTGATTTGCAGTCTAAAATGCACGAACGT',
            'f_139': 'TTTTGACAACACCTGATTTTGCAGTTAAAATGCACGAACGT',
            'f_140': 'TTTTGACAACACCTGATCTTTGCAGTAAAATGCACGAACGT',
            'f_141': 'TTTTGACAACACCTGATCTTTTGCATAAAATGCACGAACGT',
            'f_142': 'TTTTGACAACACCTGATCTGTTTGCTAAAATGCACGAACGT',
            'f_143': 'TTTTGACAACACCTGATCTGCTTTGTAAAATGCACGAACGT',
            'f_152': 'TTTTGACAACACCTGATCTGCGTCGTAAAATTTGCAGTCAT',
            'r_136': 'TTTTGACAACACCTTGACTGCAAAGTAAAATGCACGAACGT',
            'r_137': 'TTTTGACAACACCTGTGACTGCAAATAAAATGCACGAACGT',
            'r_147': 'TTTTGACAACACCTGATCTGCGTCGTAAAATCAAAGAACGT',
            'r_148': 'TTTTGACAACACCTGATCTGCGTCGTAAAATGCAAAAACGT',
            'r_149': 'TTTTGACAACACCTGATCTGCGTCGTAAAATTGCAAAACGT',
            'r_150': 'TTTTGACAACACCTGATCTGCGTCGTAAAATCTGCAAACGT',
            'r_151': 'TTTTGACAACACCTGATCTGCGTCGTAAAATACTGCAAAGT',
            'r_152': 'TTTTGACAACACCTGATCTGCGTCGTAAAATGACTGCAAAT',
            'r_153': 'TTTTGACAACACCTGATCTGCGTCGTAAAATTGACTGCAAA'}

for key, value in dict(tDNAKeys).items():
    tDNAKeys[value] = key

# %% [code]
dicts = []
sumVals = list(range(3))
for i in range(3):
    dictAll = {}
    sumVals[i] = double(sum(x for x in list(D[i].values())))
    for key, value in D[i].items():
        if len(key) > 5:
                if (key.find('TGACAGACAATCGACTGCGTAGTAAAATCCCTAAACAATTTCTACTGTTGTAGAT')>-1 and
                    key.find('TGGAGCCTTTTTTTTTTGACA')>-1 and
                    key.split('_')[1][38:44] == 'TAAAAT'):
                    gRNA = key.split('AATTTCTACTGTTGTAGAT')[1][:21]
                    tDNA = key.split('_')[1][13:54]                  
                    if (gRNA in gRNAKeys) and (tDNA in tDNAKeys):
                        bc = gRNAKeys[gRNA]+'-'+tDNAKeys[tDNA]
                        if bc in dictAll:
                            dictAll[bc] += double(value)
                        else:
                            dictAll[bc] = double(value)
                    
    dictAll = OrderedDict(sorted(dictAll.items(), key=lambda t: -t[1]))
    dicts.append(OrderedDict(dictAll))

for i in range(3):
    sumVals[i] = sum(list(dicts[i].values()))
    for key, value in dicts[i].items():
        dicts[i][key] = value/sumVals[i]

# %% [code]
def extractGrowthRate(dictK, dictSK, hselect):
    RepN = []  # No growth control
    RepF = []  # Full Growth control
    canon = 'CAGTCAGTAAAATGCAGTCA'

    for key, value in dictSK.items():
        try:
            bc1, bc2 = key.split('-')
            hdist = sum(c1 != c2 for c1, c2 in zip(gRNAKeys[bc1], gRNAKeys[bc2]))
            if hdist == hselect[0]:
                RepN.append(key)
            elif hdist == hselect[1]:
                RepF.append(key)
        except:
            pass   
    ## Find Tk, time cells grew in K-selection
    sm = 0
    km = 0
    tt = 0
    N = 0
    for k in RepN:
        try:
            sm += dictSK[k]
            km += dictK[k]
            N += 1
        except:
            pass
    Tk = -log(sm/km)

    ## Find Tk, time cells grew in SK-selection
    sp = 0
    kp = 0
    ttt = 0
    N = 0
    for k in RepF:
        try:
            sp += dictSK[k]
            kp += dictK[k]
            N += 1
        except:
            pass
    Ts = log(sp/kp)+Tk

    RepG = {}
    RepGerr = {}         
    for key, value in dictK.items():
            try:
                RepG[key] = (log(dictSK[key]/dictK[key])- log(sm/km))/(log(sp/kp) - log(sm/km))
                RepGerr[key] = sqrt(ferr(dictSK[key]*sumVals[1])**2+ferr(dictK[key]*sumVals[0])**2)/log(2)/(log(sp/kp) - log(sm/km))
            except:
                pass
    RepG = OrderedDict(sorted(RepG.items(), key=lambda t: -t[1]))        
    return RepG, RepGerr 

##########
## Mid8 Dataset
##########

RepG, RepGerr = extractGrowthRate(dicts[0], dicts[1], [16, 0])


RepE = {}
for key, value in RepG.items():
    valueRescaled = max(0.001, min(value, 0.995))
    RepE[key] = -log(1-valueRescaled)

    
RepES = OrderedDict(sorted(RepE.items(), key=lambda t: -t[1]))


RepGT, RepGTerr = extractGrowthRate(dicts[0], dicts[2], [15, 0])


# %% [code]
Data = []
Data.append(['gRNA-sequence', 'Counts-K', 'Fraction-K', 'Counts-SK', 'Fraction-SK', 'Counts-TK', 'Fraction-TK', 'GrowthRate-SK', 'GrowthRateError', 'BindingEnergy-SK', 'GrowthRate-TK'])
for key, value in dicts[0].items():
    try:
        Data.append([key, 
                     str(round(value*sumVals[0])), str(value), 
                     str(round(dicts[1][key]*sumVals[1])), str(dicts[1][key]), 
                     str(round(dicts[2][key]*sumVals[2])), str(dicts[2][key]),                      
                     str(RepG[key]), str(RepGerr[key]), str(RepES[key]), str(RepGT[key])])
    except:
        pass

f = open("../csv/MigratingTargetData.csv","w+")
for s in Data:
    f.write(','.join(s)+'\n')
f.close()  

save('../npy/MigratingTargetData.npy', Data)
```
