## Supplementary material for "Massively parallel CRISPRi assays reveal concealed thermodynamic determinants of dCas12a binding": PAM-score-Analysis.ipynb

```
# %% [code]
%pylab
from collections import OrderedDict
import json
import os
from Bio import SeqIO
from scipy.interpolate import interp1d

%matplotlib inline

def reverseComplement(seq):
    complement = {'A': 'T', 'C': 'G', 'G': 'C', 'T': 'A'}
    seq = seq.upper()
    return "".join(complement.get(base, base) for base in reversed(seq))

FILEPATH = '../fastq/'

counts = loadtxt('../error/counts.txt', delimiter= '\n', unpack = False)
logerror = loadtxt('../error/logerror.txt', delimiter = '\n', unpack = False)

#We assume that this then asymtotes to the final value - so we add:
counts = np.append(counts, [1e+07])
logerror = np.append(logerror, [0.07829763])

ferr = interp1d(counts,logerror)

# %% [code]
def importfastq(fname):
    handle1 = open(fname, "rU")
    readiter1 = SeqIO.parse(handle1, "fastq")
    BC1list = [None] * 50*10**6
    k1 = 0
    for r1 in readiter1:
        BC1list[k1] = str(r1.seq)
        k1 += 1
        if (k1 % 1000000)==0:
            print(k1)
    return BC1list[:k1]


files = os.listdir(FILEPATH)

#Finding fastq files
fastq = []
for f in files:
    if f.endswith('R1.fastq') or f.endswith('R1_001.fastq'):
        fastq.append(FILEPATH+f)

#Reading fastq files

for f1 in fastq:
    f2t = f1.split('R1')
    #Check if file exists
    if not os.path.exists(FILEPATH+f2t[0][:-1]+'_DUMB.json'):
        f2 = f2t[0]+'R2'+f2t[1]
        BC1 = importfastq(f1)
        BC2 = importfastq(f2)
        # Add counts to dictionary
        dictionary = {}
        for i in range(len(BC1)):
            guide = BC1[i]+'_'+reverseComplement(BC2[i])
            if guide in dictionary:
                dictionary[guide] += 1
            else:
                dictionary[guide] = 1
        # Order dictionary by counts        
        dictionary = OrderedDict(sorted(dictionary.items(), key=lambda t: -t[1]))
        # Save dictionary
        with open('../fastq/'+f2t[0][:-1]+'_DUMB.json', 'w') as fp:
            json.dump(dictionary, fp)

# %% [code]
N = ['G', 'C', 'A', 'T']
V = ['G', 'C', 'A']


dictGuide = {}
k = 0
for s1 in N:
    for s2 in N:
        for s3 in N:
            for s4 in ['T']:
                for s5 in ['T']:
                    for s6 in N:
                        guide = s1+s2+s3+s4+s5+s6
                        dictGuide[guide] = k
                        k += 1
                        


# %% [code]
PAMK = json.load(open('../fastq/9852_9824_79320_BY6JG_6PAM_K_TTAGGC_DUMB.json'))
PAMSK = json.load(open('../fastq/9852_9824_79322_BY6JG_6PAM_SK_CAGATC_DUMB.json'))

PAMK = OrderedDict(sorted(PAMK.items(), key=lambda t: -t[1]))
PAMSK = OrderedDict(sorted(PAMSK.items(), key=lambda t: -t[1]))

D = [PAMK, PAMSK]

# %% [code]
# Cas12-Repression
Rejects = []
dicts = []
sumVals = list(range(2))
for i in range(2):
    dictAll = {}
    for key, value in D[i].items():
        if len(key) > 5:
            try:
                if (key.find('TGACAGACAATCGACTGCGTAGTAAAATCCCTAAACAATTTCTACTGTTGTAGATCAGTCAGTAAAATGCAGTCAA')>-1 and
                    key.find('TTTTGGAGCCTTTTTTTTTTGACAACCT')>-1 and
                    key.find('CAGTCAGTAAAATGCACGAATGTTGGGGATGAAGAAATGTAC')>-1):
                    bc = key.split('CAGTCAGTAAAATGCACGAATGTTGGGGATGAAGAAATGTAC')[0][-6:]
                    if bc in dictAll:
                        dictAll[bc] += double(value)
                    else:
                        dictAll[bc] = double(value)
                else:
                    if value > 1000:
                        Rejects.append(key+'-'+str(i)+'-'+str(value))
            except:
                pass
    dictAll = OrderedDict(sorted(dictAll.items(), key=lambda t: -t[1]))
    dicts.append(OrderedDict(dictAll))

for i in range(2):
    sumVals[i] = sum(list(dict(dicts[i]).values()))
    for key, value in dicts[i].items():
        dicts[i][key] = value/sumVals[i]

# %% [code]


RepN = []  # No growth control
RepF = []  # Full Growth control

for key, value in dicts[0].items():
    try:
        if key.endswith('T'):
            RepN.append(key)
        elif key[1:] == 'TTTTA':
            RepF.append(key)
    except:
        pass
def extractGrowthRate(dictK, dictSK):
    RepN = []  # No growth control
    RepF = []  # Full Growth control
    for key, value in dictSK.items():
        try:
            if key.endswith('T'):
                RepN.append(key)
            elif key[1:-1] == 'TTTT':
                RepF.append(key)
        except:
            pass   
    ## Find Tk, time cells grew in K-selection
    sm = 0
    km = 0
    tt = 0
    N = 0
    for k in RepN:
        try:
            sm += dictSK[k]
            km += dictK[k]
            N += 1
        except:
            pass
    Tk = -log(sm/km)

    ## Find Tk, time cells grew in SK-selection
    sp = 0
    kp = 0
    ttt = 0
    N = 0
    for k in RepF:
        try:
            sp += dictSK[k]
            kp += dictK[k]
            N += 1
        except:
            pass
    Ts = log(sp/kp)+Tk

    RepG = {}
    RepGerr = {} 
    for key, value in dictK.items():
            try:
                RepG[key] = (log(dictSK[key]/dictK[key])- log(sm/km))/(log(sp/kp) - log(sm/km))
                RepGerr[key] = sqrt(ferr(dictSK[key]*sumVals[1])**2+ferr(dictK[key]*sumVals[0])**2)/log(2)/(log(sp/kp) - log(sm/km))
            except:
                pass
    RepG = OrderedDict(sorted(RepG.items(), key=lambda t: -t[1]))        
    return RepG, RepGerr  

##########
## Mid8 Dataset
##########

RepG1, RepG1err = extractGrowthRate(dicts[0], dicts[1])


RepE = {}
for key, value in RepG1.items():
    valueRescaled = max(0.001, min(value, 0.995))
    RepE[key] = -log(1-valueRescaled)

    
RepE1 = OrderedDict(sorted(RepE.items(), key=lambda t: -t[1]))


# %% [code]
Data = []
Data.append(['PAM-sequence', 'Counts-K', 'Fraction-K', 'Counts-SK', 'Fraction-SK', 'GrowthRate', 'GrowthRateError', 'BindingEnergy'])
for key, value in RepG1.items():

        Data.append([key, 
                     str(round(dicts[0][key]*sumVals[0])), str(dicts[0][key]), 
                     str(round(dicts[1][key]*sumVals[1])), str(dicts[1][key]), 
                     str(RepG1[key]), str(RepG1err[key]), str(RepE1[key])])

f = open("../csv/PAMData.csv","w+")

for s in Data:
    f.write(','.join(s)+'\n')
f.close()                                                                                 
    

save('../npy/PAMData.npy', Data)
```
