## Supplementary material for "Massively parallel CRISPRi assays reveal concealed thermodynamic determinants of dCas12a binding": Plus1Targets-Analysis.ipynb

```
# %% [code]
%pylab
from collections import OrderedDict
import json
import os
from Bio import SeqIO
from scipy.interpolate import interp1d

%matplotlib inline

def reverseComplement(seq):
    complement = {'A': 'T', 'C': 'G', 'G': 'C', 'T': 'A'}
    seq = seq.upper()
    return "".join(complement.get(base, base) for base in reversed(seq))

def Complement(seq):
    complement = {'A': 'T', 'C': 'G', 'G': 'C', 'T': 'A'}
    seq = seq.upper()
    return "".join(complement.get(base, base) for base in (seq))

FILEPATH = '../fastq/'

counts = loadtxt('../error/counts.txt', delimiter= '\n', unpack = False)
logerror = loadtxt('../error/logerror.txt', delimiter = '\n', unpack = False)

#We assume that this then asymptotes to the final value - so we add:
counts = np.append(counts, [1e+07])
logerror = np.append(logerror, [0.07829763])

ferr = interp1d(counts,logerror)

# %% [code]
def importfastq(fname):
    handle1 = open(fname, "rU")
    readiter1 = SeqIO.parse(handle1, "fastq")
    BC1list = [None] * 50*10**6
    k1 = 0
    for r1 in readiter1:
        BC1list[k1] = str(r1.seq)
        k1 += 1
        if (k1 % 1000000)==0:
            print(k1)
    return BC1list[:k1]


files = os.listdir(FILEPATH)

#Finding fastq files
fastq = []
for f in files:
    if f.endswith('R1.fastq') or f.endswith('R1_001.fastq'):
        fastq.append(FILEPATH+f)

#Reading fastq files

for f1 in fastq:
    f2t = f1.split('R1')
    #Check if file exists
    if not os.path.exists(FILEPATH+f2t[0][:-1]+'_DUMB.json'):
        f2 = f2t[0]+'R2'+f2t[1]
        BC1 = importfastq(f1)
        BC2 = importfastq(f2)
        # Add counts to dictionary
        dictionary = {}
        for i in range(len(BC1)):
            guide = BC1[i]+'_'+reverseComplement(BC2[i])
            if guide in dictionary:
                dictionary[guide] += 1
            else:
                dictionary[guide] = 1
        # Order dictionary by counts        
        dictionary = OrderedDict(sorted(dictionary.items(), key=lambda t: -t[1]))
        # Save dictionary
        with open('../fastq/'+f2t[0][:-1]+'_DUMB.json', 'w') as fp:
            json.dump(dictionary, fp)

# %% [code]
Gib = json.load(open('../fastq/Cas12a-Target-Repeat-Gibson_S2_L001_DUMB.json'))
Gib = OrderedDict(sorted(Gib.items(), key=lambda t: -t[1]))

KK = json.load(open('../fastq/Cgate-Repeat-K_S1_L001_DUMB.json'))
KK = OrderedDict(sorted(KK.items(), key=lambda t: -t[1]))
SK = json.load(open('../fastq/Cgate-Repeat-SK_S2_L001_DUMB.json'))
SK = OrderedDict(sorted(SK.items(), key=lambda t: -t[1]))

D = [Gib, KK, SK]

# %% [code]
dictGuide = {reverseComplement('AGCGTTAGCGCGTCATGGACCTTCTTGTGGACACCTAATGCCTGTAGCTGGACGCCTGAGT').upper(): 'r0',
     reverseComplement('AGCGTTAGCGCGTCATGGACCTTCTTGTGGACACCTAttcgtgcattttactgactgcaaa').upper(): 'r27',
     reverseComplement('AGCGTTAGCGCGTCATGGACCTTCTTGTGGACACCTttcgtgcattttactgactgcaaaT').upper(): 'r28',
     reverseComplement('AGCGTTAGCGCGTCATGGACCTTCTTGTGGACACCttcgtgcattttactgactgcaaaGT').upper(): 'r29',
     reverseComplement('AGCGTTAGCGCGTCATGGACCTTCTTGTGGACACttcgtgcattttactgactgcaaaAGT').upper(): 'r30',
     reverseComplement('AGCGTTAGCGCGTCATGGACCTTCTTGTGGACttcgtgcattttactgactgcaaaTGAGT').upper(): 'r32',
     reverseComplement('AGCGTTAGCGCGTCATGGACCTTCTTGTGGttcgtgcattttactgactgcaaaCCTGAGT').upper(): 'r34',
     reverseComplement('AGCGTTAGCGCGTCATGGACCTTCTTGTttcgtgcattttactgactgcaaaCGCCTGAGT').upper(): 'r36',
     reverseComplement('AGCGTTAGCGCGTCATGGACCTTCTTttcgtgcattttactgactgcaaaGACGCCTGAGT').upper(): 'r38',
     reverseComplement('AGCGTTAGCGCGTCATGGACCTTCttcgtgcattttactgactgcaaaTGGACGCCTGAGT').upper(): 'r40',
     reverseComplement('AGCGTTAGCGCGTCATGGACCTttcgtgcattttactgactgcaaaGCTGGACGCCTGAGT').upper(): 'r42',
     reverseComplement('AGCGTTAGCGCGTCATGGACttcgtgcattttactgactgcaaaTAGCTGGACGCCTGAGT').upper(): 'r44',
     reverseComplement('AGCGTTAGCGCGTCATGGttcgtgcattttactgactgcaaaTGTAGCTGGACGCCTGAGT').upper(): 'r46',
     reverseComplement('AGCGTTAGCGCGTCATttcgtgcattttactgactgcaaaCCTGTAGCTGGACGCCTGAGT').upper(): 'r48',
     reverseComplement('AGCGTTAGCGCGTCttcgtgcattttactgactgcaaaTGCCTGTAGCTGGACGCCTGAGT').upper(): 'r50',
     reverseComplement('AGCGTTAGCGCGttcgtgcattttactgactgcaaaAATGCCTGTAGCTGGACGCCTGAGT').upper(): 'r52',
     reverseComplement('AGCGTTAGCGttcgtgcattttactgactgcaaaCTAATGCCTGTAGCTGGACGCCTGAGT').upper(): 'r54',
     reverseComplement('AGCGTTAGttcgtgcattttactgactgcaaaACCTAATGCCTGTAGCTGGACGCCTGAGT').upper(): 'r56',
     reverseComplement('AGCGTTttcgtgcattttactgactgcaaaACACCTAATGCCTGTAGCTGGACGCCTGAGT').upper(): 'r58',
     reverseComplement('AGCGttcgtgcattttactgactgcaaaGGACACCTAATGCCTGTAGCTGGACGCCTGAGT').upper(): 'r60',
     reverseComplement('AGttcgtgcattttactgactgcaaaGTGGACACCTAATGCCTGTAGCTGGACGCCTGAGT').upper(): 'r62',
     reverseComplement('AttcgtgcattttactgactgcaaaTGTGGACACCTAATGCCTGTAGCTGGACGCCTGAGT').upper(): 'r63',
     reverseComplement('ttcgtgcattttactgactgcaaaTTGTGGACACCTAATGCCTGTAGCTGGACGCCTGAGT').upper(): 'r64',
     'ACCTTCAACTGCGTGACACACGGCAAGGCTATGGACGGTTCTCGGTCTGACCACCGACTCATAGT'.upper(): 'f0',
     'ACCTTCAACTGCGTGACACACGGCAAGGCTATGGACGGTTCtttgcagtcagtaaaatgcacgaa'.upper(): 'f129',
     'ACCTTCAACTGCGTGACACACGGCAAGGCTATGGACGGTtttgcagtcagtaaaatgcacgaaGT'.upper(): 'f127',
     'ACCTTCAACTGCGTGACACACGGCAAGGCTATGGACGtttgcagtcagtaaaatgcacgaaTAGT'.upper(): 'f125',
     'ACCTTCAACTGCGTGACACACGGCAAGGCTATGGAtttgcagtcagtaaaatgcacgaaCATAGT'.upper(): 'f123',
     'ACCTTCAACTGCGTGACACACGGCAAGGCTATGtttgcagtcagtaaaatgcacgaaCTCATAGT'.upper(): 'f121',
     'ACCTTCAACTGCGTGACACACGGCAAGGCTAtttgcagtcagtaaaatgcacgaaGACTCATAGT'.upper(): 'f119',
     'ACCTTCAACTGCGTGACACACGGCAAGGCtttgcagtcagtaaaatgcacgaaCCGACTCATAGT'.upper(): 'f117',
     'ACCTTCAACTGCGTGACACACGGCAAGtttgcagtcagtaaaatgcacgaaCACCGACTCATAGT'.upper(): 'f115',
     'ACCTTCAACTGCGTGACACACGGCAtttgcagtcagtaaaatgcacgaaACCACCGACTCATAGT'.upper(): 'f113',
     'ACCTTCAACTGCGTGACACACGGtttgcagtcagtaaaatgcacgaaTGACCACCGACTCATAGT'.upper(): 'f111',
     'ACCTTCAACTGCGTGACACACtttgcagtcagtaaaatgcacgaaTCTGACCACCGACTCATAGT'.upper(): 'f109',
     'ACCTTCAACTGCGTGACACtttgcagtcagtaaaatgcacgaaGGTCTGACCACCGACTCATAGT'.upper(): 'f107',
     'ACCTTCAACTGCGTGACtttgcagtcagtaaaatgcacgaaTCGGTCTGACCACCGACTCATAGT'.upper(): 'f105',
     'ACCTTCAACTGCGTGtttgcagtcagtaaaatgcacgaaTCTCGGTCTGACCACCGACTCATAGT'.upper(): 'f103',
     'ACCTTCAACTGCGtttgcagtcagtaaaatgcacgaaGTTCTCGGTCTGACCACCGACTCATAGT'.upper(): 'f101',
     'ACCTTCAACTGtttgcagtcagtaaaatgcacgaaCGGTTCTCGGTCTGACCACCGACTCATAGT'.upper(): 'f99',
     'ACCTTCAACtttgcagtcagtaaaatgcacgaaGACGGTTCTCGGTCTGACCACCGACTCATAGT'.upper(): 'f97',
     'ACCTTCAtttgcagtcagtaaaatgcacgaaTGGACGGTTCTCGGTCTGACCACCGACTCATAGT'.upper(): 'f95',
     'ACCTTtttgcagtcagtaaaatgcacgaaTATGGACGGTTCTCGGTCTGACCACCGACTCATAGT'.upper(): 'f93',
     'ACCtttgcagtcagtaaaatgcacgaaGCTATGGACGGTTCTCGGTCTGACCACCGACTCATAGT'.upper(): 'f91',
     'AtttgcagtcagtaaaatgcacgaaAGGCTATGGACGGTTCTCGGTCTGACCACCGACTCATAGT'.upper(): 'f89',
     'tttgcagtcagtaaaatgcacgaaAAGGCTATGGACGGTTCTCGGTCTGACCACCGACTCATAGT'.upper(): 'f88',}
 

# %% [code]
# Cas12-Repression
AA = []
dicts = []
sumVals = range(3)
for i in range(3):
    dictAll = {}
    for key, value in D[i].items():
        if len(key) > 5:
            if (key.find('TGACAGACAATCGACTGCGTAGTAAAATCCCTAAACAATTTCTACTGTTGTAGATCAGTCAGTAAAATGCACGAA')>-1 and
                key.find('CACGAATGTTGGGGATGAAGAAATGTAC') > -1):
                try:
                    seq = key.split('_')[1].split('GAAGAAATGTAC')[1]
                    seq2 = seq.split('TTTGCAGTCAGTAAAATGCAC')
                    if len(seq2) == 2:
                        bc = str(len(seq2[0])+4)+'-0'
                    elif len(seq2) == 3:
                        bc = str(len(seq2[0])+4)+'-'+str(len(seq2[0])+21+len(seq2[1]))
                    else:
                        AA.append([key, value])
                        bc = '0-0'
                    if bc in dictAll:  
                        dictAll[bc] += double(value)
                    else:
                        dictAll[bc] = double(value)
                except:
                    pass
    dictAll = OrderedDict(sorted(dictAll.items(), key=lambda t: -t[1]))
    dicts.append(OrderedDict(dictAll))

for i in range(1, 3):
    sumVals[i] = sum(dicts[i].values())
    for key, value in dicts[i].items():
        dicts[i][key] = value/sumVals[i]

# %% [code]
def extractGrowthRate(dictK, dictSK, num):
    RepN = []  # No growth control
    RepF = []  # Full Growth control
    RepN.append('0-0')
    for key, value in dictSK.items():
        try:
            bc1, bc2 = key.split('-')
            if num == '1':
                if bc2 == '0' and bc1 != '0':
                    RepF.append(key)
            elif num == '2':
                if bc2 != '0' and bc1 != '0':
                    RepF.append(key)                
        except:
            pass   
    ## Find Tk, time cells grew in K-selection
    sm = 0
    km = 0
    tt = 0
    N = 0
    for k in RepN:
        try:
            sm += dictSK[k]
            km += dictK[k]
            N += 1
        except:
            pass
    Tk = -log(sm/km)

    ## Find Tk, time cells grew in SK-selection
    sp = 0
    kp = 0
    ttt = 0
    N = 0
    for k in RepF:
        try:
            sp += dictSK[k]
            kp += dictK[k]
            N += 1
        except:
            pass
    Ts = log(sp/kp)+Tk

    RepG = {}
    RepGerr = {}
    for key, value in dictK.items():
            try:
                RepG[key] = (log(dictSK[key]/dictK[key])- log(sm/km))/(log(sp/kp) - log(sm/km))
                RepGerr[key] = sqrt(ferr(dictSK[key]*sumVals[2])**2+ferr(dictK[key]*sumVals[1])**2)/log(2)/(log(sp/kp) - log(sm/km))                
            except:
                pass
    RepG = OrderedDict(sorted(RepG.items(), key=lambda t: -t[1]))        
    return RepG, RepGerr  

##########
## Mid8 Dataset
##########

RepG, RepG1err = extractGrowthRate(dicts[1], dicts[2], '1')


RepE = {}
for key, value in RepG.items():
    valueRescaled = max(0.001, min(value, 0.995))
    RepE[key] = -log(1-valueRescaled)

    
RepE1 = OrderedDict(sorted(RepE.items(), key=lambda t: -t[1]))


# %% [code]
Data = []
Data.append(['gRNA-sequence', 'Counts-K', 'Fraction-K', 'Counts-SK', 'Fraction-SK', 'GrowthRate-SK', 'GrowthRate-SK-Error', 'BindingEnergy-SK'])
for key, value in dicts[1].items():
    try:
        Data.append([key, 
                     str(round(value*sumVals[1])), str(value), 
                     str(round(dicts[2][key]*sumVals[2])), str(dicts[1][key]),                     
                     str(RepG[key]), str(RepG1err[key]), str(RepE1[key])])
    except:
        pass

f = open("../csv/TargetPlus1-Data.csv","w+")
for s in Data:
    f.write(','.join(s)+'\n')
f.close()  

save('../csv/TargetPlus1-Data.npy', Data)
```
