## Supplementary material for "Massively parallel CRISPRi assays reveal concealed thermodynamic determinants of dCas12a binding": readme.txt

This archive contains all the scripts needed to process the data presented in the manuscript titled ``Massively parallel CRISPRi assays reveal concealed thermodynamic determinants of dCas12a binding''

First, download the raw sequencing data into the fastq/ folder.

A Jupyter notebook instance should be launched from the ipynb/ folder.
Each .ipynb file processes the raw data and outputs a csv file containing the raw counts and the growth rates in each condition in the csv/ folder. 
A copy of the data will also be saved as a .npy pickle file in the npy/ folder.
It may take time to generate required .json files the first time any of the ipynb is run. 
 
Dependencies.
Anaconda - conda.io 
BioPython - https://biopython.org/
